# Pseudouridine synthases modify human pre-mRNA co-transcriptionally and affect splicing

**DOI:** 10.1101/2020.08.29.273565

**Authors:** Nicole M. Martinez, Amanda Su, Julia K. Nussbacher, Margaret C. Burns, Cassandra Schaening, Shashank Sathe, Gene W. Yeo, Wendy V. Gilbert

## Abstract

Eukaryotic messenger RNAs are extensively decorated with modified nucleotides and the resulting epitranscriptome plays important regulatory roles in cells ^1^. Pseudouridine (Ψ) is a modified nucleotide that is prevalent in human mRNAs and can be dynamically regulated ^2–5^. However, it is unclear when in their life cycle RNAs become pseudouridylated and what the endogenous functions of mRNA pseudouridylation are. To determine if pseudouridine is added co-transcriptionally, we conducted pseudouridine profiling ^2^ on chromatin-associated RNA to reveal thousands of intronic pseudouridines in nascent pre-mRNA at locations that are significantly associated with alternatively spliced exons, enriched near splice sites, and overlap hundreds of binding sites for regulatory RNA binding proteins. Multiple distinct pseudouridine synthases with tissue-specific expression pseudouridylate pre-mRNA sites, and genetic manipulation of the predominant pre-mRNA modifying pseudouridine synthases PUS1, PUS7 and RPUSD4 induced widespread changes in alternative splicing in cells, supporting a role for pre-mRNA pseudouridylation in alternative splicing regulation. Consistently, we find that individual pseudouridines identified in cells are sufficient to directly affect splicing *in vitro*. Together with previously observed effects of artificial pseudouridylation on RNA-RNA^6–8^ and RNA-protein ^9–11^ interactions that are relevant for splicing, our results demonstrate widespread co-transcriptional pre-mRNA pseudouridylation and establish the enormous potential for this RNA modification to control human gene expression.

## Main

Pseudouridine is the most abundant modified nucleotide in RNA, and pseudouridylation of non-coding RNAs of the translation and splicing machineries is important for their functions. Recent transcriptome-wide methods for the detection of pseudouridine (Ψ) revealed that mRNAs in yeast and human cells contain pseudouridines that are dynamically regulated during development and in response to cellular stress ^2–5^. However, while roles for other mRNA modifications have been uncovered, such as that of N6-methyladenosine (m6A) in regulating splicing, export, translation and decay, the endogenous functions of pseudouridine in mRNA are still unknown ^1,12,13^. Because previous studies profiled pseudouridines in mature poly(A)+ mRNAs ^2–5^, it was unknown when mRNA becomes pseudouridylated and which steps of the mRNA life cycle are potentially affected by pseudouridines. In yeast, most mRNA pseudouridines have been genetically assigned to two conserved pseudouridine synthases (PUS), Pus1 and Pus7 ^2,3^, which are nuclear localized during normal growth ^14^. Likewise, human PUS enzymes that have been reported to pseudouridylate mRNA targets ^4,15,16^ localize to the nucleus, or have nuclear isoforms^15,17,18^. Further, PUS7 has been shown to be chromatin-associated and co-purifies with active Pol II promoters and enhancers^18^. Additionally, pseudouridines are present in nuclear resident ncRNAs ^2,3^. Thus, human PUS enzymes are present and active in the nucleus where they could target pre-mRNA. Because artificial RNA pseudouridylation has been shown to affect RNA-RNA^6–8^ and RNA-protein interactions ^9–11^ known, or likely to be relevant to splicing, we hypothesized that human PUS enzymes pseudouridylate nascent pre-mRNA where they could function in nuclear pre-mRNA processing, including splicing regulation.

Here, we used Pseudo-seq profiling on chromatin-associated nascent RNA to discover thousands of pseudouridines in pre-mRNA from the human hepatocellular carcinoma cell line HepG2. These pseudouridines are significantly enriched in the introns flanking sites of alternative splicing, and we identified widespread differences in alternative splicing in response to the genetic manipulation of pre-mRNA pseudouridylating enzymes PUS1, PUS7 and RPUSD4. We further showed that site-specific installation of a single endogenous pseudouridine is sufficient to directly influence splicing *in vitro*. Finally, intronic pseudouridines overlap the experimentally validated binding sites of dozens of RNA binding proteins (RBPs) interrogated by eCLIP, providing a mechanistic link to splicing regulation.

### Pre-mRNA is pseudouridylated co-transcriptionally in human cells

To determine if pseudouridine is added to pre-mRNA co-transcriptionally, we isolated chromatin-associated RNA from the hepatocellular carcinoma cell line HepG2 by a biochemical cellular fractionation that enriches for intron-containing unspliced pre-mRNA (Figure 1a) ^19,20^. The majority (74%) of pre-mRNA sequencing reads mapped to introns, and read coverage of introns and exons was relatively uniform demonstrating efficient capture of intronic reads from unspliced pre-mRNA (Figure 1a). This result contrasts with the primarily exonic reads observed in poly(A)+ mRNA (Figure 1a). We then performed Pseudo-seq ^2^ on 11 biological replicates of chromatin-associated RNA to identify high-confidence sites of reproducible pre-mRNA pseudouridylation. In Pseudo-seq, pseudouridines are selectively modified with the chemical N-cyclohexyl-N’-beta-(4-methylmorpholinium) ethylcarbodiimide p-tosylate (CMC). The bulky covalent CMC-pseudouridine adduct blocks reverse transcriptase and allows for sequencing-based detection of pseudouridines from truncated cDNAs (Figure 1b). By design, Pseudo-seq detects only modification sites where a substantial fraction of the RNA is modified: because pseudouridine-containing RNAs are not pre-enriched during the Pseudo-seq protocol, high stoichiometry pseudouridylation is required to generate a block to reverse transcription that is sufficiently penetrant to produce detectable peaks of Pseudo-seq signal ^21^, which is the difference in normalized reads from the CMC condition compared to the mock-treated control (Figure 1c). Using this approach, we identified known pseudouridine sites in ribosomal RNA (rRNA) with high sensitivity, specificity and reproducibility from the chromatin-associated RNA samples (Extended Data Fig. 1a, Supplementary Table 1).

**Figure 1.**
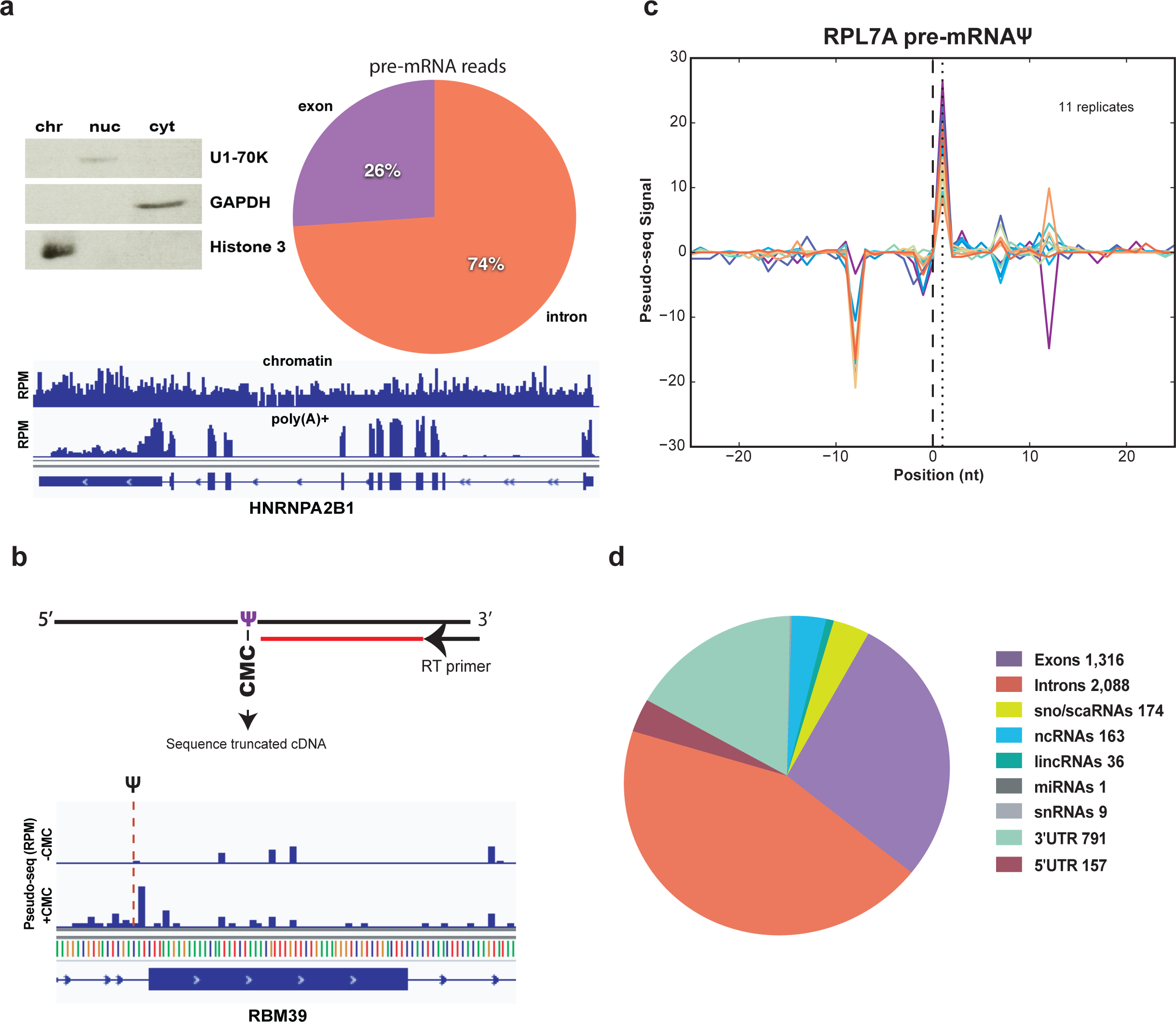
Pre-mRNA is pseudouridylated co-transcriptionally in human cells. a) Left panel. Western blot of HepG2 cellular fractions, equal cell volumes were loaded and probed with antibodies against GAPDH (cytoplasm), U1-70K (nucleoplasm) and Histone3 (chromatin). Right panel. Distribution of pre-mRNA reads mapping to introns versus exons in the chromatin-associated RNA fraction. Bottom panel. Genome browser view of reads per million mapping across the highly expressed gene hnRNPA2B1 in a chromatin-associated RNA library compared to a poly(A)+ mRNA library from HepG2 cells. b) Detection of pseudouridine by Pseudo-seq with a representative genome browser view of Pseudo-seq reads mapping to RBM39, red dotted line indicates the CMC-dependent reverse transcriptase stop corresponding to a pseudouridine (chr20:34297199). Reads per million (RPM). c) Pseudo-seq signal, equal to the difference in normalized reads between the +CMC and mock libraries. Traces for 11 biological replicates of chromatin-associated RPL7A pre-mRNA pseudouridine (chr9:136217792) are shown. d) Summary of pseudouridines identified in chromatin associated RNAs.

In addition to expected sites in rRNA (Supplementary Table 1) and snRNAs (Supplementary Table 2), analysis of Pseudo-seq signal in pre-mRNA conservatively identified thousands of novel pseudouridines, with the majority of pseudouridines found in introns (Figure 1d, Supplementary Table 3) similar to the distribution of uridines passing our read cutoff for pseudouridine detection (Extended Data Figure 2b). This number likely represents a small fraction of the total pseudouridines in pre-mRNA since only ∼1% of uridines in the nascent transcriptome had sufficient sequencing depth to meet our stringent pseudouridine calling criteria (Methods), which have generated site lists with high validation rates (>61%) in independent studies ^2,16,22^ including using completely orthologous chemistry for pseudouridine detection ^22^ (Extended Data Fig. 1b-f, Supplementary Table 4). In contrast to the apparently widespread modification of introns with pseudouridine, the only other mRNA modification that has been shown to be added to pre-mRNA, N6-methyladenosine (m6A), is found primarily in exons (90% of m6As) despite a similar distribution of intronic to exonic reads as observed in our study ^24^ (Extended Data Fig. 2a) and intronic enrichment of the motif recognized by the METTL3 enzyme that produces m6A. Hundreds more novel pseudouridines were found in nuclear resident non-coding RNAs that interact with chromatin, including the small nucleolar RNAs (snoRNAs), small cajal body-specific RNAs (scaRNAs) and long non-coding RNAs lncRNAs (Figure 1d, Supplementary Table 2, Extended Data Fig. 2c.

### Pseudouridines are enriched near alternative splice sites and RNA binding protein binding sites

This application of Pseudo-seq to study sub-cellular fractions shows that pre-mRNA is pseudouridylated in the nucleus co-transcriptionally and before splicing. To identify potential functions of pseudouridines in pre-mRNA processing, we computationally compared intronic pre-mRNA pseudouridines with RNA binding protein (RBP) binding sites identified by enhanced UV crosslinking followed by immunoprecipitation and sequencing (eCLIP-seq) for 103 RBPs from HepG2 cells ^25,26^ (Figure 2a-c). Binding sites for each examined RBP overlap tens to hundreds of pseudouridines as demonstrated by eCLIP peaks with significant enrichment over the size matched input (> 4 fold enrichment and adjusted p-value < 0.001) (Figure 2b,c). Strikingly, we find 5,359 significant eCLIP clusters (RBP binding sites) that overlap pseudouridines and 40% (1,922/4789) of unique pseudouridines overlap validated RBP binding sites across the transcriptome, of which 386 are located in introns (Supplementary Table 5). We determined the statistical significance of pseudouridine co-localization within the binding sites for individual RBPs by comparing the fraction of eCLIP peaks that overlap pseudouridines to the expected overlap for sites that were randomly located (shuffled) within intronic regions (Figure 2a). Z-scores were generated for each RBP by performing a thousand shuffles (Supplementary Table 6), revealing 33 RBPs with a Z-score above 10 (p<10^−5^) (Figure 2a). As another control, to account for the possibility of preferential detection of binding to abundant uridines, we performed the same experiment with all unmodified uridines that met the criteria for pseudouridine detection (Figure 2b). All RBPs were under-represented for intersection with only uridines, except for U2AF2, which is known to bind a uridine-rich motif (Extended Data Fig.3a).

**Figure 2.**
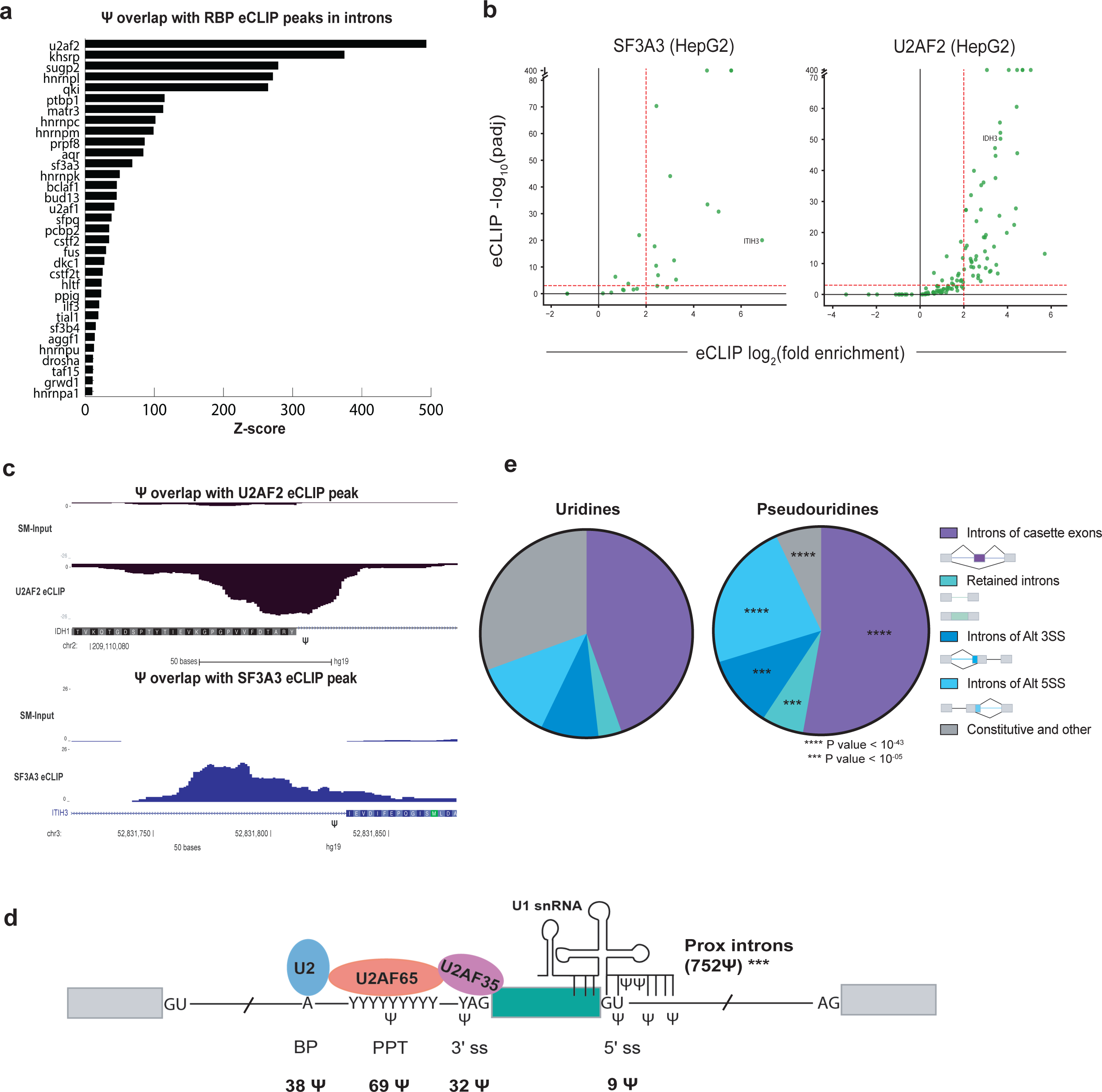
Pseudouridines are enriched near splice sites and RNA binding protein binding sites. a) Z scores were generated by comparing the fraction of eCLIP peaks overlapping pseudouridines to the calculated overlap after shuffling pseudouridines within intronic regions 1000 times. b) Volcano plots of pseudouridines overlapping RBP eCLIP peaks displaying the fold enrichment (IP over size matched input) versus the SMI-normalized adjusted *p-*value (Van Nostrand et al. 2016 Nat Methods). The overlap between eCLIP peaks and pseudouridines is shown for two RBPs U2AF2 and SF3A3. Proximal introns refer to intronic sequences <500 nt from splice sites and distal introns refer to intronic sequences >500 nt from splice sites. c) Genome browser views of U2AF2 and SF3A3 eCLIP peaks and size-matched input controls (SMI) on IDH1 and ITIH3 respectively. The location of pseudouridine relative to the eCLIP peak is denoted by (Ψ). d) Schematic of a spliced exon including the core signals for splice site recognition: branch point region (BP), polypyrimidine tract (PPT) and 5’ and 3’ splice site (ss). Number of pseudouridines identified in each splice site region is summarized below the schematic. Pseudouridines are enriched in proximal introns (within 500nt) of splice sites, *p-*value < 10^−13^ from Fisher’s exact test and within splice sites (within 6 nt from intron ends), *p*-value < 10^−6^ from Fisher’s exact test. e) Distribution of pseudouridines versus uridines with adequate read coverage in the introns of annotated alternatively spliced regions. Asterisks denote regions that are significantly enriched or depleted in the pseudouridine relative to the uridine distribution as determined by Fisher’s exact test. Alternative splice sites (Alt SS).

Many of the highest scoring RBPs with binding sites that overlap intronic pseudouridines have documented roles in splicing regulation, including core splicing factors such as U2AF2, U2AF1, SF3A3 and PRPF8 (Figure 2a-c, Supplementary Table 5). Other high scoring RBPs include polyuridine and polypyrimidine binding splicing factors such as hnRNP C, PTBP1 and TIA1 (Figure 2a, Supplementary Table 5) consistent with an enrichment for polyuridine and polypyrimidine in the sequences flanking pseudouridines (Extended Data Fig. 3b). These and other RBPs are known to be associated with nuclear RNAs and have diverse roles in RNA metabolism ^27–31^. Altogether, these results establish widespread co-occurrence of newly identified intronic pseudouridines within the binding sites of regulatory RBPs.

We found that intronic pseudouridines are enriched within splice sites (p < 10^−6^) and in proximal introns (p < 10^−13^), within 500nt of splice sites, where splicing regulatory elements such as intronic splicing enhancers and silencers are often found (Figure 2d, Supplementary Table 3). Given that the binding sites of multiple core splice site recognition factors significantly overlap sites of pseudouridylation (Figure 2a), we interrogated whether pseudouridines are present in the canonical splice site sequences. We identified pseudouridines at 3’ (32) and 5’ (9) splice sites, polypyrimidine tracts (PPT) (72) and branch site regions (37) (Figure 2d). Pseudouridines occur at critical residues for splicing including the conserved pyrimidine before the 3’ splice site (<u>Y</u>AG), in the region of the 5’ splice site that base pairs with U1 snRNA, and in the branch site region that base pairs with U2 snRNA. Our identification of pseudouridines enriched in the binding sites of diverse splicing factors and RBPs, in proximal introns, at splice sites, and within pre-mRNAs encoding splicing factors (Extended Data Fig. 3c, Supplementary Table 7) demonstrate the vast potential of pseudouridines to control nuclear pre-mRNA processing.

We further explored the potential for co-transcriptional pre-mRNA pseudouridylation to regulate splicing by determining the distribution of intronic pseudouridines with respect to alternatively spliced regions. Pseudouridines are notably enriched around alternative splice sites (p<10^−05^) including in the introns flanking cassette exons, introns of alternative 5’ and 3’ splice sites and in retained introns (Figure 2e). In contrast, pseudouridines are significantly depleted (p<10^−42^) from the introns of constitutive and other exons (Figure 2e). Together, the distribution of pseudouridines in alternatively spliced regions, near splice sites, and within splicing factor binding sites is consistent with a role for pseudouridine in splicing regulation by pre-mRNA modifying pseudouridine synthases (PUSs).

### PUS1, RPUSD4 and PUS7 are predominant pre-mRNA modifying enzymes

To identify which pseudouridine synthases (PUSs) act on pre-mRNA and potentially regulate splicing, we took an in vitro approach. Accurate pseudouridine assignment in cells requires very high sequencing depth of PUS-depleted cells in order to interpret an absence of reads as evidence of modification by that PUS. A recently developed high-throughput in vitro pseudouridylation assay ^16^ overcomes this limitation to identify which PUS(s) directly pseudouridylate sites of interest, including in lowly expressed RNAs. We verified the excellent agreement between genetic and *in vitro* assignment approaches by cross-validating 85% of yeast *PUS1*-dependent pseudouridine sites in mRNA ^2,16^ (Extended Data Fig. 4a). To identify human PUSs that directly pseudouridylate pre-mRNA sequences, we synthesized a pool of RNA containing each pseudouridine site flanked by 130 nucleotides of endogenous sequence (Figure 3a). Pseudouridylation of the RNA from the pool was detected by Pseudo-seq following incubation with individual recombinant human pseudouridine synthases PUS1, PUS7, PUS7L, PUS10, RPUSD2, RPUSD4, TRUB1, TRUB2 or HepG2 nuclear extract. Each of the 8 tested enzymes pseudouridylated specific pre-mRNA sequences in both exons and introns (Figure 3b, c, Supplementary Table 6). Some additional sites were only pseudouridylated in nuclear extract implying that at least one additional PUS also pseudouridylates pre-mRNA and/or that some sites require co-factors that are supplied in the nuclear extract.

**Figure 3.**
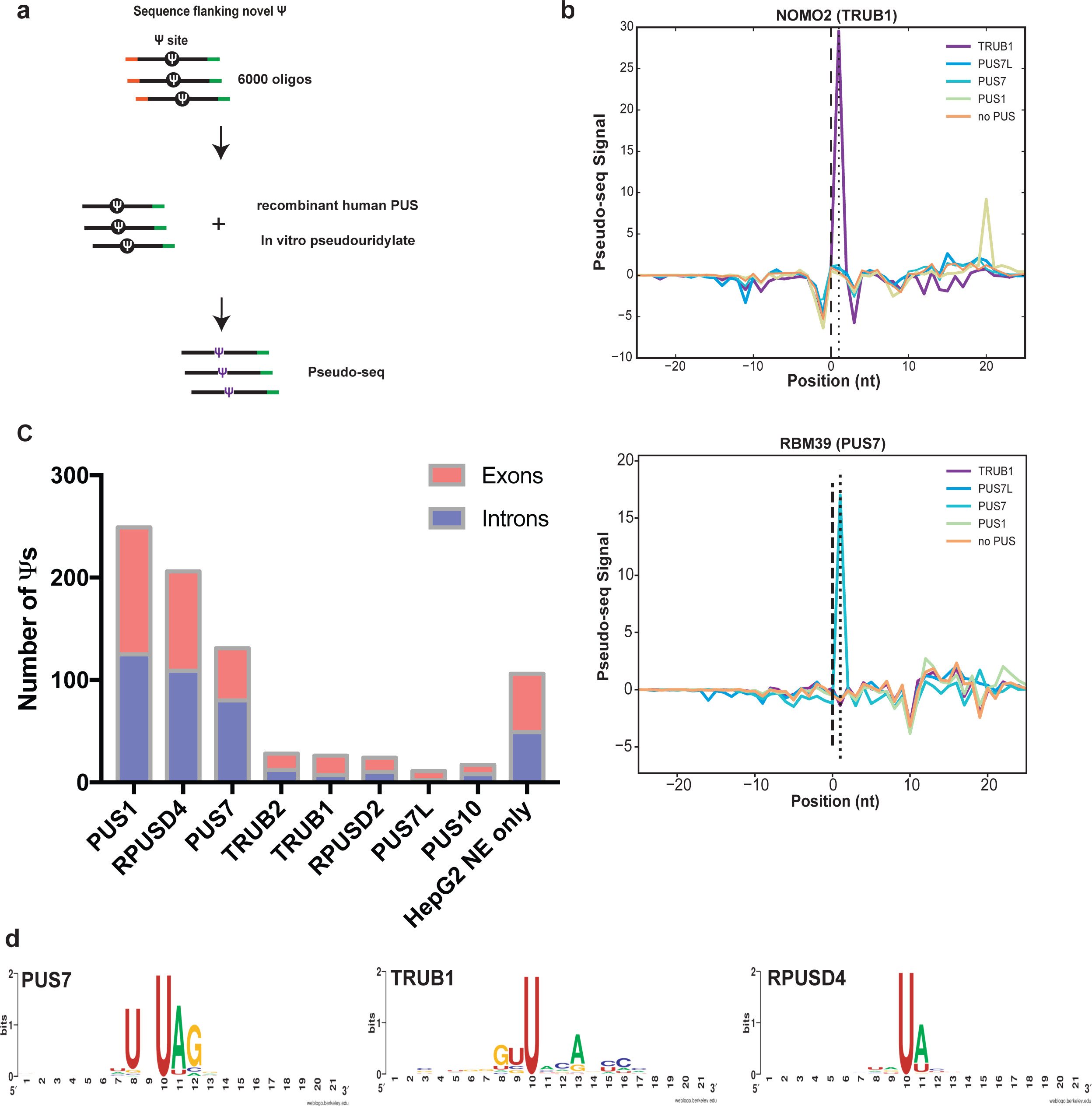
Multiple PUS pseudouridylate pre-mRNA sequences. a) Schematic of in vitro pseudouridylation assay with RNA made from a pool of 6000 oligos containing all the sites identified in HepG2 chromatin-associated RNA. In vitro pseudouridylation was carried by incubating pool RNA with recombinant human pseudouridine synthases (PUS) and pseudouridines were identified by Pseudo-seq. b) Pseudo-seq signal plots of two intronic pre-mRNA pseudouridines: a TRUB1 target in NOMO2 and a PUS7 target in RBM39. c) Summary of pseudouridines in introns and exons assigned as direct targets of each PUS protein from *in vitro* Pseudo-seq assay. d) Weblogo summarizing frequency of motifs identified among targets of PUS7, TRUB1 and RPUSD4.

We identified PUS1, RPUSD4 and PUS7 as predominant pre-mRNA pseudouridylating enzymes (Figure 3c). Orthologous yeast proteins Pus1 and Pus7 pseudouridylate the majority of known yeast mRNA sites ^2,3^, suggesting broad conservation of mRNA targeting by these PUS. RPUSD4 is a member of a PUS family that has expanded to include 4 paralogs in higher eukaryotes (chordates) and was not previously known to have mRNA targets. Importantly, PUS1, PUS7 and RPUSD4 are present in the nucleus in human cells^17,18,32^ where they have access to pre-mRNA. This nuclear function of RPUSD4 is distinct from its reported role in regulating mitochondrial 16S rRNA ^33,34^. There is almost no redundancy among the *in vitro* targets of each PUS (Extended Data Fig. 4b-d), suggesting distinct gene regulatory programs. The fact that PUS1, RPUSD4 and PUS7 pseudouridylate the majority of pre-mRNA sites is consistent with their relatively high expression in HepG2 cells (Extended Data Fig. 5a). Notably, the expression levels of the PUS proteins vary across tissues and cell types (Extended Data Fig. 5b,c) highlighting the regulatory potential of pre-mRNA pseudouridylation by these factors.

Most PUS proteins pseudouridylate their targets in diverse sequence contexts and do not display strong sequence preferences (Extended Data Fig. 4e). This may reflect a predominantly structural mode of mRNA target recognition as shown for yeast Pus1 ^16^. By contrast, the targets of PUS7 are highly enriched for a UNΨAR motif, which is similar to, but a more permissive version of the motif recognized by yeast Pus7 (Figure 3d). This motif overlaps with the YAG of the 3’ splice site for multiple PUS7 targets (Supplementary Table 8). The PUS7 motif is enriched among all chromatin-associated pseudouridines identified in HepG2 cells (Extended Data Fig. 4f) including some sites that are not modified by the recombinant protein in vitro. A failure to be pseudouridylated in vitro could reflect the need for additional RNA sequence, cellular co-factors, or other features present in the endogenous context such as co-transcriptional PUS recruitment. TRUB1 modified comparatively few targets (27) (Figure 3c), consistent with the fact that only 10 pre-mRNA pseudouridines occur in the preferred TRUB1 sequence context GUΨYNANNC ^15^ (Extended Data Fig. 3e,f). Some sites that lack the motif are nevertheless efficiently modified by TRUB1 *in vitro* revealing some flexibility in target recognition by this PUS (Figure 3d). Together, our *in vivo* pseudouridine profiling and *in vitro* PUS target validation identify pre-mRNA sites as the largest class of PUS targets and reveal the potential for multiple pseudouridine synthases to influence pre-mRNA processing by pseudouridylating diverse nascent pre-mRNA sequences.

### Pseudouridine synthases PUS1, RPUSD4 and PUS7 regulate alternative splicing

The prevalence of pseudouridines in alternatively spliced pre-mRNAs (Figure 2e) and within splicing factor binding sites (Figure 2a-d), together with evidence that pseudouridine affects diverse RNA-RNA ^6–8^ and RNA-protein ^9–11^ interactions, suggests that altering PUS activity could cause changes in splicing. PUS1, PUS7 and RPUSD4 emerged as likely candidates to influence splicing from our *in vitro* pre-mRNA pseudouridylation assay (Figure 3c). We first examined whether PUS1-dependent pseudouridylation influenced splicing by making PUS1 knockout HepG2 cells using CRISPR/CAS9 (Figure 4a). We obtained highly reproducible RNA-seq data from poly(A)+ mRNA from PUS1 knockout and wildtype cells (R^2^>0.96, Extended Data Figure 6a), and quantified differences in alternative splicing and mRNA abundance (Methods). Strikingly, PUS1 knockout leads to thousands of changes in alternative splicing, as defined by splicing events that were statistically significant (FDR < 0.05) and exhibited greater than 10% difference in inclusion levels compared to wildtype cells (Figure 4a, Supplementary Table 9). These changes in splicing affected 1617 genes and included cassette exons, alternative 5’ and 3’ splice sites, retained introns and mutually exclusive exons. We observed both differential inclusion and skipping of cassette exons upon PUS1 knockout (Figure 4b, Supplementary Table 9), showing that the effect of PUS1 on splicing is pre-mRNA dependent. In contrast to these broad effects on alternative splicing, mRNA abundance changed very little, with approximately one hundred mRNAs significantly altered in cells lacking PUS1 (Extended Data Fig. 8a). The affected mRNAs do not include splicing factors or RBPs that could indirectly affect splicing (Supplementary Table 10).

**Figure 4.**
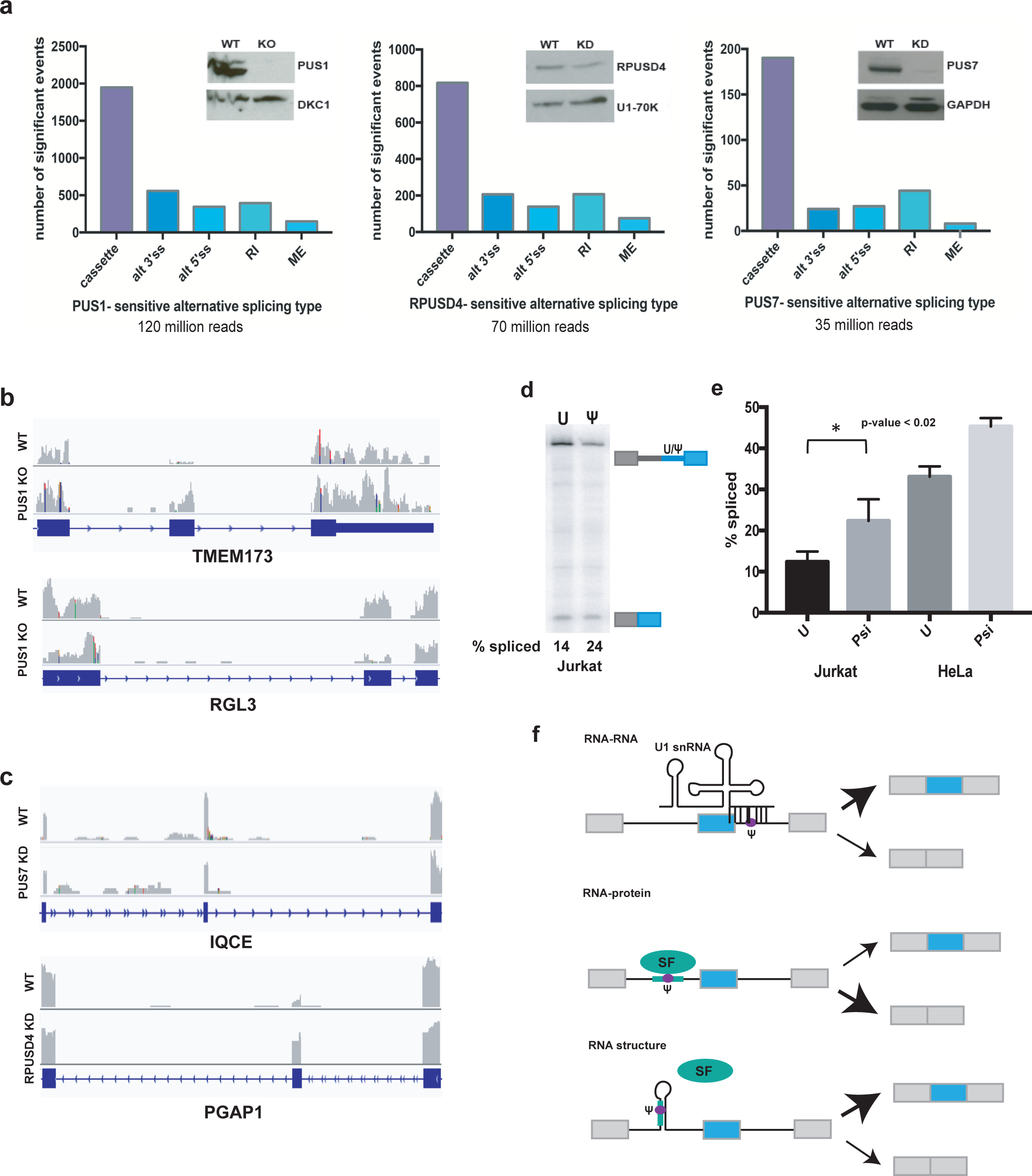
Pseudouridine synthases regulate alternative splicing and pseudouridines directly affect splicing. a) Left - Western blot of the CRISPR knockout PUS1 HepG2 cell line probed for PUS1 and a loading control. RNA was isolated from PUS1 knockout (KO) and wild type (WT) cells and mRNA-seq libraries were prepared from poly(A)+ mRNA. The number of significant alternative splicing changes in PUS1 KO versus WT (n=2 biological replicates) is displayed by type of alternative splicing: cassette exons (cassette), alternative 3’ splice sites (alt 3’ss), alternative 5’ splice sites (alt 5’ ss), retained introns (RI) and mutually exclusive exons (ME). Significant alternative splicing events were determined from rMATS as those events that changed by greater than 10% difference in percent inclusion and a false discovery rate (FDR) of less than or equal to 0.05. Middle - Western blot of representative RPUSD4 knockdown (∼60%) at 96h following shRNA induction. RPUSD4-sensitive alternative splicing changes determined from RNA-seq analysis (n=2 biological replicates) as above. Right - Western blot of representative PUS7 knockdown (∼90%) at 96h following shRNA induction. PUS7-sensitive alternative splicing changes determined from RNA-seq analysis (n=3 biological replicates) as above. b) Genome browser view of two PUS1-sensitive alternative splicing events: a cassette exon in TMEM173 that is included and in RGL3 that is skipped upon PUS1 KO. c) Genome browser view of a PUS7 enhanced cassette exon in IQCE and an RPUSD4 repressed cassette exon in PGAP1. d) Representative RT-PCR gel of *in vitro* splicing of RBM39 two-exon reporter (Extended Data Figure 8a) (–/+) a PUS7-dependent Ψ in splicing competent Jurkat nuclear extract. e) Quantification of *in vitro* splicing of the RBM39 reporter (–/+ Ψ) in Jurkat and HeLa nuclear extract. Quantification of % spliced from n=3 is displayed. f) Potential mechanisms of pseudouridine-sensitive alternative splicing.

In budding yeast, a single pseudouridine Ψ44 in the U2 snRNA is modified by Pus1. However, the corresponding position in the human U2 snRNA, Ψ43, was not affected in PUS1 KO cells (Extended Data Fig. 7a). This result is consistent with a previous report of PUS1-independent pseudouridylation of this U2 snRNA site in mouse PUS1 KO cells ^35^ and with computational and biochemical evidence that human snRNAs are modified by the snoRNA/scaRNA-dependent pseudouridine synthase, DKC1 ^36,37^. Similarly, none of the other detected U2 snRNA Ψs were affected in PUS1 KO HepG2 cells (Extended Data Fig. 7a,b) or mouse cells ^35^. These results show that PUS1-sensitive alternative splicing events are not caused by loss of U2 snRNA pseudouridylation. Consistent with the possibility of direct effects of PUS1-dependent pre-mRNA pseudouridylation on splicing, *in vitro* validated PUS1 targets in cassette exons or flanking introns of ANKRD10, PUM2, SNHG12 and TANK showed differential splicing in PUS1 KO cells (Extended Data Fig. 7c). Because pre-mRNA pseudouridine discovery was necessarily restricted to highly expressed genes, whereas orders of magnitude fewer reads are sufficient to quantify alternative splicing in mature poly(A)+ mRNA, we were unable to interrogate pseudouridine status for the vast majority of PUS1-sensitive cassette exons and flanking introns (Extended Data Fig. 7d, Supplementary Table 9).

We used inducible shRNA expression to deplete the essential ^38,39^ pre-mRNA pseudouridylating enzyme RPUSD4 and also PUS7 to determine their impact on alternative splicing by RNA-seq. Replicate experiments were reproducible (R^2^>0.98 and R^2^>0.94, Extended Data Fig. 6b,c). Partial depletion of RPUSD4 (60 %) and PUS7 (90%) produced widespread effects on alternative splicing including 817 RPUSD4-sensitive cassette exons in 696 genes and 190 PUS7-sensitive cassette exons in 175 genes (Figure 4a,c, Supplementary Table 11,12). The RPUSD4 and PUS7 samples were sequenced at lower depth than the PUS1 KO cells possibly contributing to fewer identified splicing changes. As with PUS1, we were unable to interrogate pseudouridine status of most RPUSD4- and PUS7-sensitive exons (Extended Data Figure 7d). PUS7 depletion resulted in 374 differentially expressed genes (Extended Data Figure 8b, Supplementary Table 13), consistent with previous studies showing altered PUS7-dependent mRNA levels in yeast ^3^. In contrast, RPUSD4 depletion resulted in almost no significant changes in mRNA levels (Extended Data Figure 8c, Supplementary Table 14). As expected for shRNA mediated depletion, PUS7 and RPUSD4 mRNAs were significantly downregulated (Extended Data Fig. 8b,c). Depletion of each of the predominant pre-mRNA targeting PUS produced distinct changes in the pattern of alternative splicing (Extended Data Fig. 7e). Notably, the pervasiveness and magnitude of PUS-dependent splicing changes are comparable to the effects of depleting canonical splicing regulators ^40^. These results support a widespread and non-redundant role for multiple PUSs in alternative splicing regulation. Given the enrichment of pseudouridines around alternatively spliced regions, we expect a subset of the splicing changes in PUS knockout/knockdown cells to be a consequence of direct pre-mRNA pseudouridylation, but we cannot rule out that some of the PUS-sensitive events are an indirect consequence of reduced pseudouridylation of other targets.

### Site-specific pseudouridylation directly affects splicing *in vitro*

We took advantage of our ability to site-specifically pseudouridylate pre-mRNA sequences *in vitro* (Figure 3) to determine if individual Ψs identified in cells are sufficient to directly affect splicing. We generated chimeric two-exon pre-mRNA splicing reporters containing intronic PUS7 target sites in RBM39 and MDM2 (Extended Data Fig. 9a) that were pseudouridylated in vitro with purified PUS7 (Figure 3b, Extended Data Fig. 9b) and incubated with nuclear extract under splicing conditions. Remarkably, a single endogenous intronic PUS7-dependent Ψ upstream of the 3’ splice site in the RBM39 pre-mRNA was sufficient to directly enhance splicing as quantified by RT-PCR and compared to splicing of the unmodified control (Figure 4d,e). This effect of Ψ on splicing was observed in nuclear extract from two different cell lines (Figure 4e, Extended Data Fig. 9c) and across time points (Extended Data Fig. 9d). Similarly, an MDM2 splicing reporter with a Ψ at a 3’ splice site UAG directly enhanced splicing *in vitro* (Extended Data Fig. 9e,f). These *in vitro* experiments avoid all potentially confounding indirect effects of genetic manipulation of PUS activity in cells and the results demonstrate that there is a direct mechanistic effect of individual endogenous pre-mRNA pseudouridines on splicing.

## Discussion

Our study of chromatin-associated RNA revealed wide-spread co-transcriptional modification of pre-mRNA with pseudouridine at more than four thousand locations, positioning pseudouridine to influence virtually all steps of mRNA processing. Pre-mRNA pseudouridylation is likely to be even more extensive than the sites identified here because stringent criteria for detection and the large size of the nascent transcriptome restricted intronic pseudouridine discovery to highly expressed genes. About half of the newly identified pseudouridines are located in introns, where they were not previously known to occur. These pseudouridines are well positioned to affect alternative splicing due to their enrichment in proximal introns, alternatively spliced regions and within splicing factor binding sites. Consistent with this regulatory potential, we find that installation of a single intronic pseudouridine is sufficient to affect splicing outcome in vitro, and genetic manipulation of pre-mRNA pseudouridylating enzymes in cells leads to thousands of changes in alternative splicing.

Mechanistically, pseudouridines in pre-mRNA have the potential to influence splicing by three main mechanisms: altering pre-mRNA-snRNA interactions, modulating pre-mRNA-protein interactions, or by influencing pre-mRNA secondary structure (Figure 4f) ^13^. We identified pseudouridines in regions poised to function by any of these modes. Single pseudouridines stabilize synthetic RNA duplexes by 1-2 kcal/mol as compared to uridine base pairs ^6,7^. This stabilizing effect is predicted to enhance splice site recognition by promoting binding of spliceosomal snRNAs to the pseudouridylated 5’ splice site or branch site. Pseudouridines have also been shown to alter the affinities of various RNA binding proteins (e.g. PUM2, MBNL1, U2AF65) for RNA ^9–11^. We find that approximately 25% of intronic pseudouridines are located within experimentally validated binding sites for 84 RBPs analyzed in the same cell line, which is significantly more than expected by chance (Figure 2d-f). The sensitivity of most RBPs to pseudouridylation of their binding sites remains to be determined, but many are likely to be affected given that diverse RBPs show 2 to 100-fold differences in affinity for pseudouridylated compared to unmodified RNA ^9–11^. Finally, pseudouridylation could indirectly alter splice site accessibility and/or splicing factor binding by changing pre-mRNA secondary structure, as has recently demonstrated for certain intronic m6A modification sites ^41^. Altogether, our results implicate pseudouridine and the pre-mRNA modifying pseudouridine synthases as novel regulators of pre-mRNA processing. This may have clinical relevance for the multiple pseudouridine synthases implicated in mitochondrial myopathy ^17,42^, digestive disorders ^43–47^, intellectual disability ^48,49^, resistance to viral infection ^50,51^, X-linked dyskeratosis congenita^52,53^, and cancer ^54–57^.

## Methods

### Cell Culture

Human hepatocellular carcinoma cells HepG2 from ATCC HB8065 (lot 59635738) were grown in DMEM (HyClone SH30022.FS) supplemented with 10% fetal bovine serum (FBS HyClone SH30071.03). Cells were grown at 37C with 5% CO2 and maintained at subconfluency.

### Total RNA Isolation

HepG2 cells were harvested by pelleting and resuspending fresh or frozen (−80C) pellets in 1mL of QIAzol (Qiagen). Total RNA was harvested according to the manufacturer’s procedure.

### Western Blotting

Whole cell lysates were made by pelleting HepG2 cells and re-suspending fresh or frozen (−80C) pellets in RIPA buffer (50mM Tris pH 8, 150 mM NaCl, sodium deoxycholate 0.5%, sodium dodecyl sulfate 0.1%, NP-40 1%), lysed on ice for 10 min with vortexing. Spun down at 4C and maximum speed (13,200 rpm) for 15 min and collected supernatant as lysate. Approximately, 20ug of whole cell lysates, as determined by Bradford assay, were loaded in 10% SDS-PAGE for western blot of PUS1 knockout and wildtype HepG2 cells. HepG2 fractions were isolated as described below and treated with Benzonuclease to release proteins from nucleic acid. Equal cell volumes of cellular fractions (3%) were loaded in 12% SDS-PAGE gels for Western blots of cell compartments to determine fraction purity. Gels were transferred to nitrocellulose membranes. Membranes were blocked in 5% milk and incubated with primary antibodies overnight at 4C in 5% milk low-salt TBST (50 mM Tris pH 7.5 150 mM NaCl 0.1% Tween-20). Antibodies used for Western blot were as follows: anti-PUS1 at 1:1000 (Bethyl Labs A301-651A), anti-U1-70K at 1:2000 (Millipore 05-1588), anti-GAPDH at 1:10,000 (Sigma-Aldrich G9545), anti-H3 at 1:20,000 and anti-DKC1 at 1:1000 (GeneTex GTX109000). Secondary antibody incubation was for 1 hour at room temperature using HRP conjugated antibodies: anti-mouse IgG at 1:3000(Invitrogen 62–6520) or goat anti-rabbit IgG at 1:3000 (Promega W4011). Washes were with high-salt TBST (50 mM Tris pH 7.5 400 mM NaCl 0.1% Tween-20).

### CRISPR knockout generation

PUS1 CRISPR knockout HepG2 cells were generated using a two-guide strategy to take out the first exon containing the translation start codon. Oligos for the upstream PUS1upF 5’-CACCGCGCAGGGTCCACCGTCCGA -3’ and PUS1upR 5’-AAACTCGGACGGTGGACCCTGCGC -3’, and for the downstream guide PUS1dnF 5’-CACCGATAACAGCGGTTAGCGGCA -3’ and PUS1dnR 5’-AAACTGCCGCTAACCGCTGTTATC -3’ were phosphorylated and annealed and then cloned into px458 (Addgene) digested with BbsI. HepG2 cells were transfected with both plasmids for the upstream guide and downstream guide (1.25 µg of each) with Lipofectamine 2000 according to the manufacturer’s protocol in 6-well plates. After 24h the cells were visually inspected for GFP fluorescence and prepared for sorting by trypsinizing and re-suspended in FACS buffer (PBS without Mg2+ and Ca2+ and supplemented with 2% FBS). Single GFP positive cells were sorted on an Aria I sorter and plated into each well of a 96 well plate. PUS1 knockout clones were expanded and knockout verified by PCR of genomic DNA and Western Blot with anti-PUS1.

### PUS protein depletion

shRNAs targeting PUS7 and RPUSD4 were cloned into the lentiviral vector pLKO-Tet-On (Addgene) digested with AgeI-HF and EcoRI-HF to remove the stuffer sequence. Two oligos containing the complementary shRNA targeting sequence with the corresponding overhangs were annealed and ligated into the vector by standard cloning. Oligos sequences for RPUSD4 and PUS7 were:

shRPUSD4_2F

5’-CCGGGCTTCGAGTTCACTTGTCCTTCTCGAGAAGGACAAGTGAACTCGAAGCTTTTTG-3’, shRPUSD4_2R, 5’-AATTCAAAAAGCTTCGAGTTCACTTGTCCTTCTCGAGAAGGACAAGTGAACTCGAAGC-3’

shPUS7L_4F

5’-CCGGTCTTAGTTCAGACTCATATATCTCGAGATATATGAGTCTGAACTAAGATTTTTG-3’

shPUS7L_4R

5’-AATTCAAAAATCTTAGTTCAGACTCATATATCTCGAGATATATGAGTCTGAACTAAGA-3’.

Lentiviral particles were prepared by transfecting a 10cm dish of HEK293T cells with 5µg pLKO-Tet-On, 4.5µg pCMV-dR8.2, 500ng pCMV-VSV-G and transfection reagent X-tremeGENE 9 according to the manufacturer’s protopcol. Viral supernatant was harvested 48h after transfection. HepG2 cells were transduced in 6-well plate with 1mL of viral supernatant in 6 well plates with 1mL of cell suspension. Cells stably integrated with the lentiviral vector were selected 48h post-transduction with 3µg/mL of puromycin until cells in the untransduced cells did not survive. shRNA expression was induced in HepG2 cells with Doxocycline to a final concentration of 500 ng/mL. Cells were maintained in Doxoxycline containing media for 96h and whole cell lysates were prepared for Western Blot analysis of depletion and RNA was isolated for RNA-seq library construction.

### Nuclear extract preparation

HepG2 cells were pelleted by spinning down at 1000 rpm for 5 min, washed with PBS. Cells were transferred to 1.5 mL tube and centrifuged at 3000 rpm for 1 minute. The pellet was re-suspended in cytoplasmic extract buffer (10 mM HEPES pH 7.6, 1.5 mM MgCl2, 10 mM KCl, 0.15% NP-40) ∼100 µL per 2×10^6^ cells and incubated on ice for 5 minutes and spun down for 3 min at 6500 rpm for 3 minutes. Supernatant was collected as cytoplasmic fraction. The nuclear pellet was re-suspended in equal volume of nuclear extract buffer (20 mM HEPES pH 7.6, 1.5 mM MgCl_2_, 420 mM NaCl, 0.2 mM EDTA, 20% glycerol). Three cycles of 15 minutes at −80C followed by thawing at 37C with vortexing for 1 minute in between cycles. Spun down at max speed for 15 minutes at 4C. Collected supernatant as nuclear extract.

### Cellular Fractionation

Biochemical fractionation was performed essentially as described in ^19,58^ for 11 biological replicates of HepG2 cells. All fractions were prepared from fresh cell pellets. Two 10 cm dishes with ∼10×10^6^ each of HepG2 cells were trypsinized and spun down at 500xg and washed with cold PBS (1mM EDTA). Cell pellets were re-suspended in 400 µL of cytoplasmic NP-40 lysis buffer (10 mM Tris-HCl pH 7.5, 0.15% NP-40, 150 mM NaCl) by flicking tube and incubated on ice for 5 minutes. Lysate was layered over 1 mL of sucrose cushion (24% RNAse-free sucrose in cytoplasmic lysis buffer) and spun down 10 minutes at 15,000xg at 4°C. Supernatant was collected as cytoplasmic fraction. Nuclear pellet was washed 2x with PBS without displacing pellet (1mM EDTA) and re-suspended in 200 µL glycerol buffer (20 mM Tris-HCl pH 7.9, 75 mM NaCl, 0.5 mM EDTA, 0.85 mM DTT, 0.125 mM PMSF, 50% glycerol) by flicking tube followed by addition of 200 µL of nuclei lysis buffer (10 mM HEPES pH 7.6, 1 mM DTT, 7.5 mM MgCl_2_, 0.2 mM EDTA, 0.3 M NaCl, 1 M UREA, 1% NP-40). Samples were then vortexed on high 2x 2 seconds, incubated on ice for 2 minutes, centrifuged for 2 min at 4°C 15,000xg and the supernatant collected as nucleoplasmic fraction. The remaining chromatin pellet was washed 2x with PBS without displacing pellet (1mM EDTA) and re-suspended in 100uL of PBS (1mM EDTA). DNase I (2uL NEB) was added to re-suspended chromatin-pellet and incubated at 37°C for 5-10 minutes to dissolve pellet. Collected 10uL for western blot of fractions and added 1mL of QIAzol (Qiagen) to remaining chromatin and incubated at 50°C for 10 min to solubilize chromatin. Followed manufacturer’s instruction to finish isolating chromatin-associated RNA with QIAzol. Samples were DNase I treated (NEB) treated after QIAzol isolation according to manufacturer’s instructions.

### rRNA depletion

Ribosomal RNA (rRNA) was depleted using one Ribozero (Illumina MRZH11124) reaction per 20ug of chromatin-associated RNA following the manufacturer’s protocol.

### Recombinant PUS plasmids and purification

Human TRUB1, PUS7, PUS7L, TRUB2, RPUSD2 and PUS10 were cloned from human cDNA obtained from Human ORFeome (cDNA from Mammalian Gene Collection) with Gibson assembly into the BamH1 site of pET15b expression. Recombinant hPus1 was purified as described ^59^. Expression was induced in BL21 (DE3) Gold cells [Agilent] with 0.1 mM IPTG at OD600 0.6-0.8. Cells were grown overnight at 16°C, then harvested by centrifugation and resuspended in lysis buffer (50 mM HEPES-KOH pH 7.0, 500 mM NaCl, 5 mM β-mercaptoethanol, 1x Protease Inhibitor Cocktail (Roche)) and lysed by sonication. Human PUS1 was induced and the bacterial lysate was centrifuged at 12,000 rpm for 30 min, bound on a HisTrap column (GE Healthcare) and eluted off the column with 250 mM imidazole. The protein was then dialyzed overnight at 4°C into storage buffer (50 mM HEPES-KOH pH 7.0, 100 mM NaCl, 1 mM β-mercaptoethanol) and further purified by gel filtration over a Superdex-200 column (GE Healthcare). The protein product was concentrated with a centrifugal filter unit (MD Millipore) and concentration determined by Bradford staining against a BSA standard. Human TRUB1, PUS7, PUS7L, TRUB2, RPUSD2 and PUS10 were purified as follows. Rosetta 2 BL21 (DE3) pLysS cells were transformed with the expression vector and an individual colony was grown at 37°C in LB to OD600 0.8. Induction of expression was overnight at 18°C with 1mM IPTG. Protein was affinity purified using the HisTrap HP 5mL column (GE) on an FPLC. Bound protein was washed with wash buffer (50mM potassium phosphate buffer pH 8, 0.5M NaCl, 30mM Imidazole) and then eluted with elution buffer (50mM potassium phosphate buffer pH 8, 0.5M NaCl, 300mM Imidazole). Protein was concentrated with Amicon Ultra Centrifugal Filter Units and stored in storage buffer containing 20mM HEPES (pH 7.5), 200mM NaCl, 10% glycerol, and 1mM DTT. Concentration was determined by Bradford with BSA standards.

### In vitro pseudouridylation

A pool of 6000 DNA oligos (Twist Biosciences) was designed to contain all the sites identified as pseudouridines in chromatin-associated RNA from HepG2 cells. The DNA oligos in the pool contained 65 nucleotides upstream and 64 nucleotides downstream of the pseudouridine. In addition, the sequences contained the T7 promoter and a 3’ adapter sequence to serve as handles for PCR amplification. The pool was amplified by PCR with primers oBZ131 (GCTAATACGACTCACTATAGGG) and oTC_pool2_rev (GTCCTTGGTGCCCGAGTG) using Phusion Polymerase (NEB). PCR reactions were supplemented with 3% DMSO and gel purified prior to in vitro transcription. RNA was in vitro transcribed with the MEGAshortscript T7 transcription kit (Thermo Fisher) and gel purified. RNA was re-suspended in water, denatured at 75C for 2 min, cooled on ice for 5 min and folded at 37C for 20 min following addition of 5X pseudouridylation buffer (500 mM Tris pH 8.0, 500 mM Ammonium Acetate, 25 mM MgCl2, 0.5 mM EDTA) to 1X for the final reaction volume. In vitro pseudouridylation reactions were carried out by incubating 15-30 pmol of folded RNA with 600nM recombinant human pseudouridine synthases: PUS1, PUS7, PUS7L PUS10, RPUSD2, RPUSD4, TRUB1, TRUB2, HepG2 nuclear extract or no PUS for 45 minutes at 30C in 500uL final reaction volumes extract in (1X pseudouridylation buffer, 2mM DTT). RNA was purified immediately by phenol chloroform extractions and isopropanol precipitated.

### Pseudo-seq

Pseudo-seq libraries were prepared as previously described in detail ^21^. Briefly, rRNA-depleted chromatin-associated RNA was fragmented in 10mM ZnOAc for 2 minutes at 60C. Fragmentation was quenched by addition of EDTA to 20mM. RNA was then either treated with CMC (0.4M final) or mock treated (−CMC) in BEU buffer for 45 min at 40C and CMC was reversed from Us and Gs by incubation in Sodium carbonate buffer for 2 hours at 50C. RNA fragments 120-140 nucleotides in length were size selected from denaturing polyacrylamide gels for library preparation. The ends of the fragmented RNA were repaired by treatment with T4 Polynucleotide Kinase (NEB) and Calf intestinal alkaline phosphatase (NEB) in 1X PNK buffer (NEB). A 3’ adenylated adapter (AppTGGAATTCTCGGGTGCCAAGG/3ddC/) was ligated to the RNA fragments with T4 RNA ligase in 1X T4 RNA ligase buffer (NEB), followed by reverse transcription with AMV RT (Promega) and RT primer: /5Phos/GATCGTCGGACTGTAGAACTCTGAACCTGTCGGTGGTCGCCGTATCATT/iSp18 /CACTCA/iSp18/GCCTTGGCACCCGAGAATTCCA. Truncated cDNAs (120-170nt) were size selected from denaturing polyacrylamide gels and gel purified cDNAs were circularized with CircLigase ssDNA ligase (Epicentre). Circularized cDNAs were PCR amplified with forward primer (AATGATACGGCGACCACCGA) and BC reverse primer (CAAGCAGAAGACGGCATACGAGATXXXXXXGTGACTGGAGTTCCTTGGCACCCGAGAATTCCA where Xs represent the unique barcode sequence). In vitro Pseudo-seq libraries were prepared as described above except with full length in vitro transcribed RNA recovered from in vitro pseudouridylation assays. After CMC modification and reversal, full length RNA was gel purified in denaturing polyacrylamide gels. Reverse transcription was performed using the 3’ adapter sequence appended to the pool oligos as a handle with RT primer ONM_RT-L2 (/5Phos/NNNNNNNNNGATCGTCGGACTGTAGAA CTCTGAACGTGTAGATC/iSp18/CACTCA/iSp18/CCTTGGCACCCGAGAATTCCAGTCCTTGGTGCCCGAGTG), truncated cDNAs from 140-170 nts were size selected and circularized for PCR amplification with primers RP1 (AATGATACGGCGACCACCGAGATCTACACGTTCAGAGTTCTACAGTCCGA) and BC reverse primer. Libraries were sequenced on an Illumina HiSeq in single-end mode to 20-30 million reads per sample for in vivo Pseudo-seq and 15-20 million reads for in vitro Pseudo-seq libraries.

/5Phos/GATCGTCGGACTGTAGAACTCTG

AACCTGTCGGTGGTCGCCGTATCATT/iSp18/CACTCA/iSp18/GCCTT

GGCACCCGAGAATTCCA

/5Phos/GATCGTCGGACTGTAGAACTCTG

AACCTGTCGGTGGTCGCCGTATCATT/iSp18/CACTCA/iSp18/GCCTT

GGCACCCGAGAATTCC

### RNA-seq

Total RNA was isolated for two replicates of wildtype and PUS1 knockout HepG2 cell as described above and stranded poly(A)+ selected mRNA-seq libraries were performed by the MIT BioMicroCenter using the TruSeq Stranded mRNA Library Prep Kit from Illumina according to the manufacturers’ protocol. Libraries were sequenced on an Illumina Next-seq with paired end 40-bp reads to a depth of ∼150 million paired reads per replicate. mRNA purity was comparable across libraries with 1.4% (WT1), 1.7% (WT2), 1.3% (KO1) and 1.3% (KO2) rRNA contamination. PUS7 (n=3) and RPUSD4 (n-2) knockdown stranded poly(A)+ selected mRNA-seq libraries were prepared by Genewiz and sequenced on a HiSeq with paired end 150-bp reads.

### Sequencing Analysis

All sequencing data has been deposited in GEO, accession GSE123613. mRNA-seq reads were mapped to the human genome using STAR aligner (version). Reads were mapped to genome assembly GRCh37 with ENSEMBL GRCh37.75 annotations. STAR alignment was carried out using the following parameters: --outFilterType BySJout --outFilterMultimapNmax 20 --alignSJoverhangMin 8 --alignSJDBoverhangMin 1 --outFilterMismatchNmax 999 --alignIntronMin 20 --alignIntronMax 1000000 --alignMatesGapMax 1000000 --alignEndsType EndToEnd. Pseudo-seq sequencing data was analyzed using in house Bash and Python scripts. Cutadapt ^60^ was used to trim the 3’ adapter sequence from the reads. In vitro Pseudo-seq sequencing reads were also PCR duplicate collapsed using fastx_collapser (http://hannonlab.cshl.edu/fastx_toolkit/). Processed reads were mapped to a bowtie index of ENSEMBL GRCh37.75, 45S rRNA and snRNAs for in vivo Pseudo-seq libraries and the pool of sequences for in vitro Pseudo-seq libraries using tophat2 ^61^. The mapped reads were processed with in house Python scripts. Pseudo-seq signal or peak height was calculated as follows for all possible used annotated in GRCh37.75 excluding repetitive elements. For each U position centered on a 51-nucleotide window, the fraction of reads whose 5’ends map to the position was divided by the reads mapping to the window was calculated. Pseudo-seq signal is the difference in the fraction of reads between the +CMC and −CMC multiplied by the window size. For each Ψ the Pseudo-seq signal corresponds to that of the expected RT stop 1 nt 3’ of the Ψ site. Sites were called as pseudouridines if an RT stop with a peak height >1, met a read cutoff (reads/nt in the window) of 0.1 and was present in 7 out of 11 replicate libraries of chromatin-associated RNA.

### PUS protein assignment

Assignment of human PUS to pseudouridylated targets was carried out by using the Grubbs outlier test with significance level alpha set to 0.05 to identify sites that had peak height values that deviated from the normal distribution of peak heights for all the other conditions (all other PUS and no PUS). A target site was assigned to a PUS if it was called as an outlier exclusively in the corresponding PUS sample and had a peak height greater than > 1.0. For nuclear extract assignments, all sites assigned to any recombinant PUS were removed and the Grubbs outlier test was performed on the remaining sites to identify sites that are pseudouridylated only in nuclear extract.

### Alternative splicing analysis

Analysis of differential alternative splicing between wildtype and PUS1 KO HepG2 mRNA-seq was carried out by rMATS (version 3.) using Ensembl GRCh37.72 annotations. rMATS reported differences in alternative splicing of types skipped exons (SE), alternative 3’ splice sites (A3SS), alternative 5’ splice sites (A5SS), retained introns (RI) and mutually exclusive exons. Percent inclusion differences were determined from the rMATS junction only output files. Events with an absolute difference in percent inclusion between PUS1 KO and WT cells of greater than or equal to 10% and with and false discovery rate (FDR) of equal to or less than 0.05 are considered significant and reported.

### Differential gene expression analysis

HTseq (version 0.6.1) was used to generate read counts tables that were submitted to DESeq2 (version 1.14.1) to determine differences in mRNA abundance between PUS1 KO, RPUSD4 KD or PUS7 KD and WT HepG2 cells from mRNA-seq data. After DESeq analysis genes with 0 counts in at least one sample were filtered out. Differences in mRNA levels with Padj < 0.05 were considered significant.

### Enrichment Analysis

Bedtools was used for overlapping pseudouridines and uridines with genomic features. As a background set for calculating the enrichment of pseudouridines in different regions we used all the uridines that met our read cutoff of 0.1 in our replicate cutoff of 7 out of 11, as was used for pseudouridine calling. Detected pseudouridines were classified into features according to gencode v19 annotations according to the following priority snoRNAs > lincRNAs > snRNA > miRNAs > antisense ncRNAs > 5’ UTR > 3’ UTR > exons > introns. Retained introns were filtered as introns. Enrichments of pseudouridines in the introns of alternatively spliced regions was carried out by overlapping intronic pseudouridines compared to uridines with MISO version 2 annotations (https://miso.readthedocs.io/en/fastmiso/annotation.html) which were generated from Ensembl genes, known Genes (UCSC) and RefSeq genes annotations. Fisher’s exact test was used in contingency tables of uridine versus pseudouridine distributions in the different features, p-values were corrected for multiple hypothesis testing using Bonferroni corrections and p-values < 0.05 were considered significant. Pseudouridines were considered as present in branch site regions if they overlapped annotated branch site regions ^62–64^. Pseudouridines were classified as present in putative PPT tracts if they were within 50 nucleotides of the 3’ splice site AG and by manual inspection.

### Motif Enrichment

131 nucleotides of sequence surrounding pseudouridine was used as the input to MEME version 5.0.2 in discriminative mode to identify motifs enriched in pseudouridine compared to the 131 nucleotides of sequence flanking the background set of uridines that met our read cutoff for pseudouridine calling in the Pseudo-seq libraries. Significantly enriched motifs are presented.

### GO analysis

Gene ontology analysis was performed for pseudouridine containing genes using PANTHER version 11 ^65^. All Ensembl gene IDs for pseudouridine containing genes were used as the test set and the Ensembl gene IDs for all genes that contained uridines matching our read cutoff for pseudouridine identification (Supplementary Table 4). Reported enriched GO terms correspond to those that were significant after Bonferroni correction for multiple hypothesis testing.

### Analysis of RBP eCLIP overlap with pseudouridines

Size matched input-normalized bed files for eCLIP biological replicates were combined into a single bedtool of shared peaks using bedtools intersect, where a shared peak was defined as at least one intersecting nucleotide. eCLIP peaks at pseudouridines in HepG2 cells were then identified by using bedtools intersect to determine eCLIP peaks where the peak overlapped with a pseudouridine. Volcano plots of these pseudouridine intersecting eCLIP peaks were then generated using the eCLIP log_2_(fold change) and Padj ^26^ values. We selected significance cutoffs of log_2_(fold change) of 2 and –log_10_(Padj) of 3. For the volcano plots, if an RBP had multiple pseudouridines within an eCLIP peak, we plotted the best cluster as defined by first the lowest Padj and highest fold change values. Introns were determined by the genic location using gencode hg19 annotations. To identify pseudouridine-interacting RBPs we ranked RBPs for preferential binding to pseudouridine sites. We calculated the fraction of significant eCLIP peaks that intersected pseudouridine sites over the total number of significant eCLIP peaks, then compared this ratio to one calculated using pseudouridine sites shuffled within introns. We performed this calculation with 1000 iterations of shuffling the pseudouridine location in introns to calculate a standard distribution of this ratio, and then calculated the z-score of the observed ratio. To filter for RBPs that bind uridine-rich motifs, we performed the same calculation with all high-confidence uridines that did not have a pseudouridine and repeated the 1000 iterations of shuffling and measuring the intersection with the RBPs.

### *In vitro* splicing from nuclear extracts

The splicing reporter backbone pcAT7-Glo1 was digested with NdeI and BglII and Gblocks including the entire endogenous exon and 100nt of intronic sequence (including the pseudouridylated position) and 15nt of vector overlapping sequence. The Gblock sequences are:

RBM39

TAGAAACTGGGCATATGATTTGAATGAACCCTGCTATTGTAGTCCTCTTTTATTAATGCTTTCCTGACATTTACCCTGTTAGTTGAGGCTCTTCATTGTTCCTGCACTGAGCTGTAGAATTCTCTTTTGTTATAGATTTGCAAACAAGACTTTCCCAGCAGACTGAAGTCTAGAGGGCCCGTTT

MDM2

TAGAAACTGGGCATATGGCTTTAGTTTTAACTGTTGTTTATGTTCTTTATATATGATGTATTTTCCACAGATGTTTCATGATTTCCAGTTTTCATCGTGTCttttttttCCTTGTAGGCAAATGTGCAATACCAACATGTCTGTACCTACTGATGGTGCTGTAACCACCTCACAGATTCCAGCTTCGGAACAAGAGACCCTGTCTAGAGGGCCCGTTT.

Plasmids were *in vitro* transcribed and subsequently linearized by restriction digest with XbaI. Splicing reporter RNA (30pmol) were mock treated for unmodified or *in vitro* pseudouridylated with recombinant human PUS7 as described in the *in vitro* pseudouridylation section. Purified RNA substrates (15 pmol) were incubated with 32% Jurkat or HeLa nuclear extract in a total volume of 13 uL under splicing conditions 11.5 mM Tris-HCl, pH7.5, 3.0 mM MgCl2, 1 mM ATP, 20 mM CP, 0.5 mM DTT, 58 mM KCl, 3% PVA, 0.1 mM EDTA, and 11.5% glycerol. Reactions were incubated for 30-60 min at 30°C or indicated time points. RNA was recovered from the reactions by proteinase K treatment, phenol-chloroform extraction and ethanol precipitation. The RNAs from the splicing reactions were analyzed by RT-PCR with reporter specific primers GE1-F – 5’-GCAAGGTGAACGTGGATGAAGTTGG – 3’ and MDM2_E-R 5’-CAGGGTCTCTTGTTCCGAAGCTGG -3’ or RBM39_E-R 5’-CAGTCTGCTGGGAAAGTCTTGTTTGC -3’.

### Site lists and reproducibility analyses

We obtained lists of pseudouridine sites called in two publications: the initial list from our group, published in supplemental table 8 of Carlile *et al.* 2014, and those reported in supplemental table S7 of Khoddami *et al.* 2019. For Extended Data Figure 1B, we compared the lists reported by either publication and categorized them into sets, based on whether they had been called by either or both of these methods. For Extended Data Figure 1C, we additionally determined the sites that had been called in any of the following publications: (a) Schwartz 2014, which called sites transcriptome wide in HEK293 cells, rather than HeLa; (b) Carlile 2019, which used an in vitro approach to validate sites (c) Li 2015. We then used the UpSetR package (version 1.4.0) to visualize the reproducibility of the Carlile 2014 sites across these datasets.

### Sequence context analyses

For each set of sites in Extended Data Figure 1B, we generated a fasta file that contained the 11-nt sequence surrounding the pseudouridine site. We then used Weblogo 3 to visualize the frequency of each nucleotide relative to the pseudouridine site.

### RBS-Seq signal analysis

We downloaded the raw reads from GSE90963, using reads obtained from RiboMinus-treated total RNA (GSM2418439 and GSM2418440) and polyA-selected RNA (GSM2418443 and GSM2418444). Adapters were trimmed using bbduk. Reads were mapped to hg19 using novoalign, using parameters -t 60 -h 120 -b 4 -H 20 -r A 2 -s 2. For bisulfite-treated samples, the-b 4 parameter was also included. Python scripts using the pysam package were used to determine coverage, the number of deletions, and the number of read 5’ ends at each position, which were then stored as wig files for visualization.

## Extended Data Figure Legends

**Extended Data Figure 1**

a) ROC curve of true positive versus false positive rates of known pseudouridine locations in human rRNA for two representative chromatin-associated RNA replicates. The +CMC and - CMC traces are displayed for both replicates. b) Validation rate of Pseudo-Seq sites in HeLa. Here we plot how many of the sites reported in Carlile *et al*, 2014 were also detected in at least one of four other approaches. The circles next to the method names indicate that a set of sites was detected in that method. Horizontal bars indicate the number of Pseudo-Seq sites that were tested by each method. c) Two chemically independent methods detect pseudouridine sites in distinct sequence contexts, suggesting capture biases. For each set of sites (sites unique to either dataset, or sites shared by both datasets), sequence logos are centered at the pseudouridine site, with the size indicating the relative frequency of each nucleotide at each position relative to the pseudouridine. d-f) Genome browser views illustrating the Pseudo-Seq signal at sample sites, with bars indicating the number of read 5’ ends at each position. These include: d) Two sites detected by Pseudo-Seq and chemically validated as direct targets of a PUS, but not called by RBS-Seq or another method; e) Highly reproducible Pseudo-Seq sites that were not detected by other techniques; f) Examples of cell-type specific pseudouridylation.

**Extended Data Figure 2**

a) Read and pseudouridine distribution in chromatin-associated RNA from HepG2 cells in this study compared to read and m6A distribution in chromatin-associated RNA from another study by ^24^. b) Percent of uridines or pseudouridines meeting the minimum read cutoff in introns compared to exons. c) Pseudo-seq signal plot of a reproducible pseudouridine on the nuclear resident ncRNA NEAT1. Data for 11 replicates of chromatin-associated RNA are shown.

**Extended Data Figure 3**

a) The z score of pseudouridines shuffled 1000 times within intronic regions plotted against the z score of uridines shuffled 1000 times within intronic regions. b) Motifs enriched around pseudouridines as determined by MEME in discriminative mode. The target sequences submitted for motif enrichment were 65 and 64 nucleotides flanking on both sides of all the pseudouridines or all background uridines detected in chromatin-associated Pseudo-seq. c) Gene ontology analysis of pseudouridine containing pre-mRNAs using PANTHER. All uridines meeting our read and replicate cutoff for pseudouridine detection were used as the background set.

**Extended Data Figure 4**

a) Validation rate of mRNA pseudouridines genetically assigned to yeast Pus1 by *in vitro* Pseudo-seq with recombinant yeast Pus1 (Carlile et al. 2019). b-d) The union of Pseudo-seq signal of all sites *in vitro* pseudouridylated with PUS1, PUS7 or RPUSD4 with Pseudo-seq signal >2 for at least one of these PUS is displayed. Target sites assigned to each PUS are colored blue (PUS1), purple (PUS7) or orange (RPUSD4). Scatter plot of Pseudo-seq signal at sites pseudouridylated by b) PUS1 compared to PUS7, c) PUS1 compared to RPUSD4 and d) PUS7 compared to RPUSD4. e) Weblogo summarizing frequency of motifs identified among in vitro targets of PUS1, PUS7, PUS7L PUS10, RPUSD2, RPUSD4, TRUB1 and TRUB2. f) Enrichment of PUS7 and TRUB1 motif among all chromatin-associated pseudouridines. The analysis was centered on the pseudouridine position and motif enrichment was calculate using CentriMO.

**Extended Data Figure 5**

a) Expression levels of all PUS protein genes in HepG2 cells derived from HepG2 WT mRNA-seq. Read counts were normalized by mRNA length for longest isoform obtained from ENSEMBL Biomart to calculate RPKM values for each. b,c) Expression levels of all PUS protein genes across a panel of human cell lines (b) and human tissues (c) expressed as TPMs and obtained from EMBL-EBI Expression Atlas and Illumina Body Map.

**Extended Data Figure 6**

Scatter plot of normalized RNA-seq read counts showing reproducibility for replicates of a) wildtype and PUS1 KO HepG2 cells, b) wildtype and PUS7 KD HepG2 cells c) wild type and wildtype and RPUSD4 KD HepG2 cells. Pearson R^2^ values among replicates are shown.

**Extended Data Figure 7**

a) Primer extension of U2 snRNA with unmodified or CMC treated total RNA from wildtype or PUS1 KO HepG2 cells using U2snRNA_prox R primer. U2 snRNA pseudouridines identified from CMC-dependent stops are labeled. Ψ43 is quantified relative to background non-pseudouridine signal in the lane below. b) Primer extension of U2 snRNA with unmodified or CMC treated total RNA from wildtype or PUS1 KO HepG2 cells using U2snRNA_dist R primer. U2 snRNA pseudouridines identified from CMC-dependent stops are labeled. c) Change in percent inclusion of PUS1-sensitive cassette exons that are targets of PUS1 *in vitro.* d) Number of PUS1, RPUSD4 or PUS7-sensitive cassette exons, where the exon and flanking intronic sequence that had <10%, >10% or >50% coverage of Us by Pseudo-seq. e) Venn diagram of overlap among cassette exons regulated by PUS1, PUS7 and RPUSD4.

**Extended Data Figure 8**

a) Volcano plot of differential expression log2 fold change and -Log10 of P*adj.* in PUS1 KO compared to WT determined by DEseq2 analysis of mRNA-seq. Changes with an adjusted *p*-value < 0.05 were considered significant. b) Volcano plot of differential expression log2 fold change and -Log10 of P*adj.* in PUS7 KD compared to WT determined by DEseq2 analysis of mRNA-seq. Changes with an adjusted *p*-value < 0.05 were considered significant. c) Volcano plot of differential expression log2 fold change and -Log10 of P*adj.* in RPUSD4 KD compared to WT determined by DEseq2 analysis of mRNA-seq. Changes with an adjusted *p*-value < 0.05 were considered significant.

**Extended Data Figure 9**

a) Schematic of two exon chimeric splicing reporters for RBM39 and MDM2. These reported include the respective exon and 100nt of upstream intronic sequence containing the pseudouridine (Ψ). Sequences in grey are derived from B-globin and in blue are from the endogenous pre-mRNA sequence. 25nt of the intron and 10nt of the exon are displayed for context. b) In vitro Pseudo-seq signal of the PUS7 installed MDM2 pseudouridine. c) Representative RT-PCR gel of in vitro splicing of the RBM39 reporter in Jurkat and HeLa nuclear extract. Quantification of RT-PCR results (n=3) is shown. d) *In vitro* splicing of the RBM39 in a splicing timecourse (30, 60 and 90 minutes) in Jurkat nuclear extract. e) In vitro splicing of the MDM2 reporter in Jurkat nuclear extract. Replicates are shown independently to reflect differences in overall splicing efficiency among replicates. All statistically significant differences are denoted by asterisks and p-values listed.

**Supplementary Table 1**

HepG2 45S rRNA pseudouridines identified in chromatin-associated RNA. Pseudo-seq signal is listed for each replicate.

**Supplementary Table 2**

HepG2 ncRNA pseudouridines identified in chromatin-associated RNA. Pseudo-seq signal is listed for each replicate.

**Supplementary Table 3**

HepG2 pre-mRNA pseudouridines identified in chromatin-associated RNA. Pseudo-seq signal is listed for each replicate.

**Supplementary Table 4**

HeLa pseudouridine validation analysis (Cassandra)

**Supplementary Table 5**

RBP eCLIP peaks with significant enrichment over the size matched input (> 4-fold enrichment and adjusted p-value < 0.001) and overlapping pseudouridine locations.

**Supplementary Table 6**

Z scores comparing the fraction of RBP eCLIP peaks overlapping pseudouridines to the calculated overlap after shuffling pseudouridines within intronic regions 1000 times.

**Supplementary Table 7**

Genes containing pseudouridine versus background set of genes that met uridine read cutoff that were used for gene ontology analysis

**Supplementary Table 8**

In vitro pseudouridylated sites by recombinant human PUSs. Pseudo-seq signal is listed for each source of PUS. PUS assignment was determined by Grubbs outlier analysis.

**Supplementary Table 9**

Cassette exons or skipped exons (SE) that are alternatively spliced in PUS1 KO compared to WT HepG2 cells as determined by rMATS.

**Supplementary Table 10**

Differentially expressed genes in PUS1 KO compared to WT HepG2 cells as determined by DESeq2 analysis of mRNA levels.

**Supplementary Table 11**

Cassette exons or skipped exons (SE) that are alternatively spliced in RPUSD4 KD compared to WT HepG2 cells as determined by rMATS.

**Supplementary Table 12**

Cassette exons or skipped exons (SE) that are alternatively spliced in PUS7 KD compared to WT HepG2 cells as determined by rMATS.

**Supplementary Table 13**

Differentially expressed genes in PUS7 KD compared to WT HepG2 cells as determined by DESeq2 analysis of mRNA levels.

**Supplementary Table 14**

Differentially expressed genes in RPUSD4 KD compared to WT HepG2 cells as determined by DESeq2 analysis of mRNA levels.

## Acknowledgements

We thank Brenton R. Graveley for HepG2 cells. We thank Karla M. Neugebauer and members of the Gilbert lab for helpful discussions and critical reading of the manuscript. We thank Tristan A. Bell for the batch of purified hPUS1. Splicing competent nuclear extracts and splicing reporter backbone plasmids were a kind gift from Matthew G Thompson and Kristen W. Lynch. This work was supported by NIH GM101316 to W.V.G; HG004659, HG009889, HL137223, HL137219 to G.W.Y. and American Cancer Society RSG-13-396-01-RMC to W.V.G., Jane Coffin Childs Post-doctoral Fellowship 161624T to N.M.M.

