## Supplemental Figures for "Pseudouridine synthases modify human pre-mRNA co-transcriptionally and affect splicing"

#### Extended Data Fig. 1

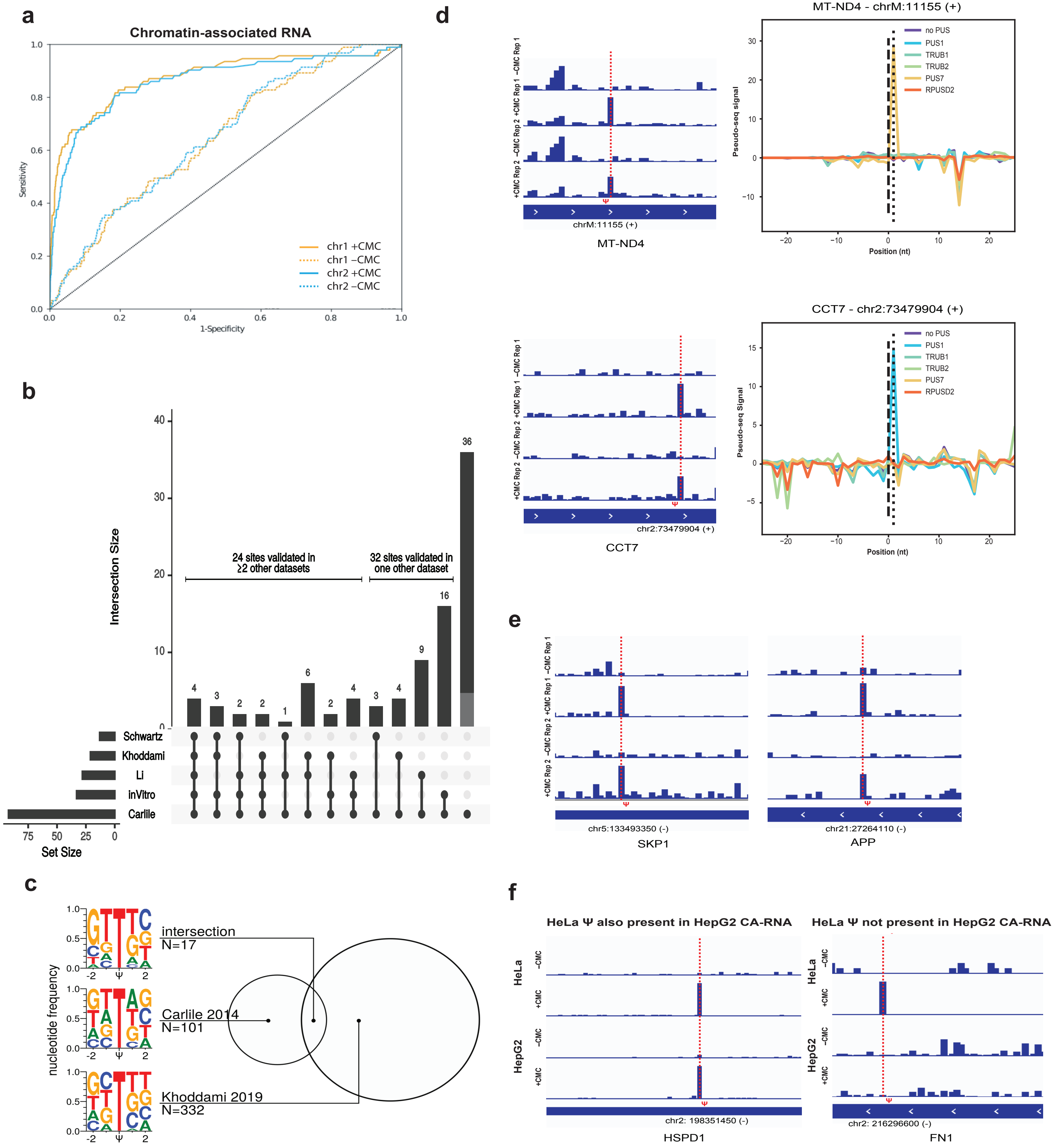

#### Extended Data Fig. 2

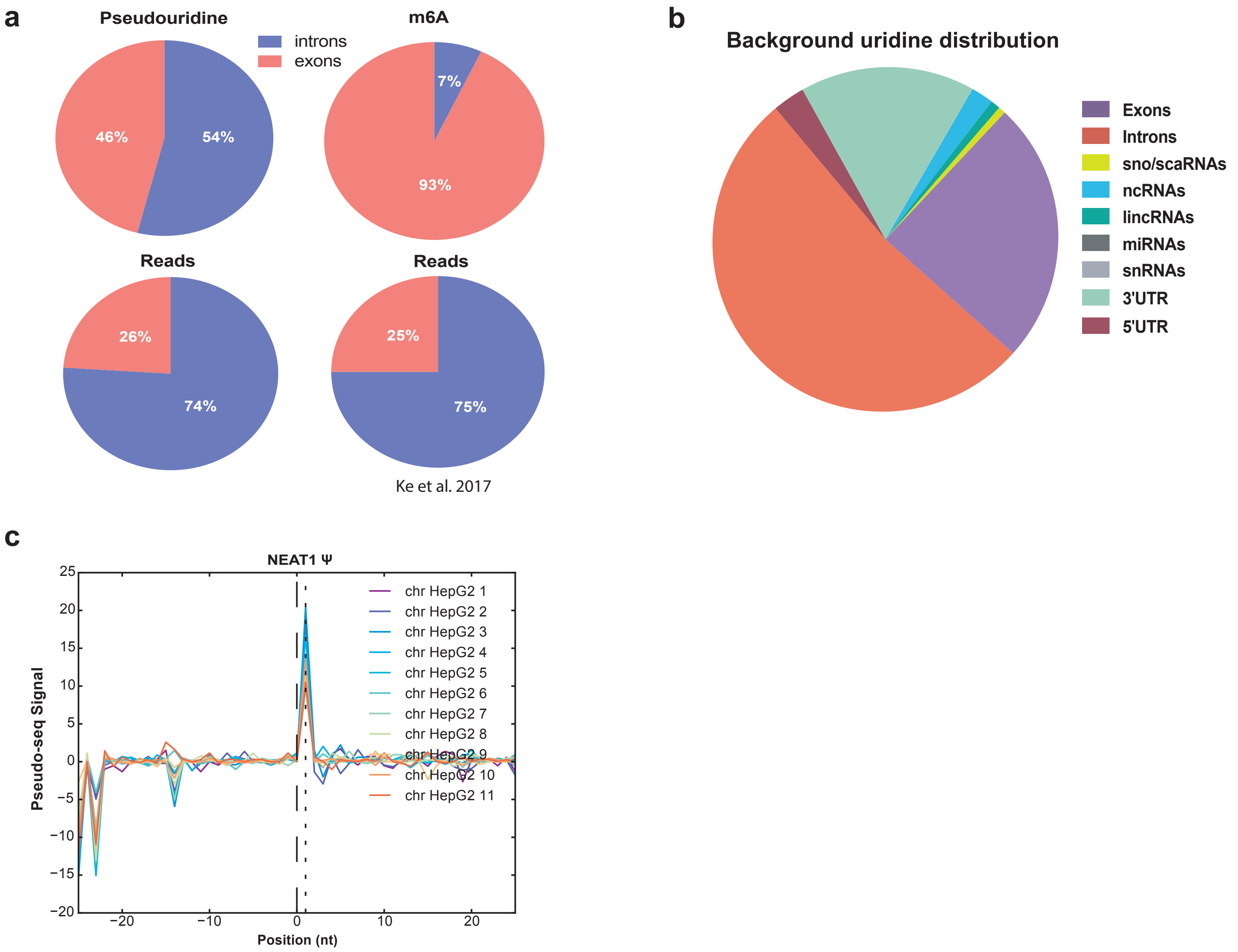

**Extended Data Fig. 3**

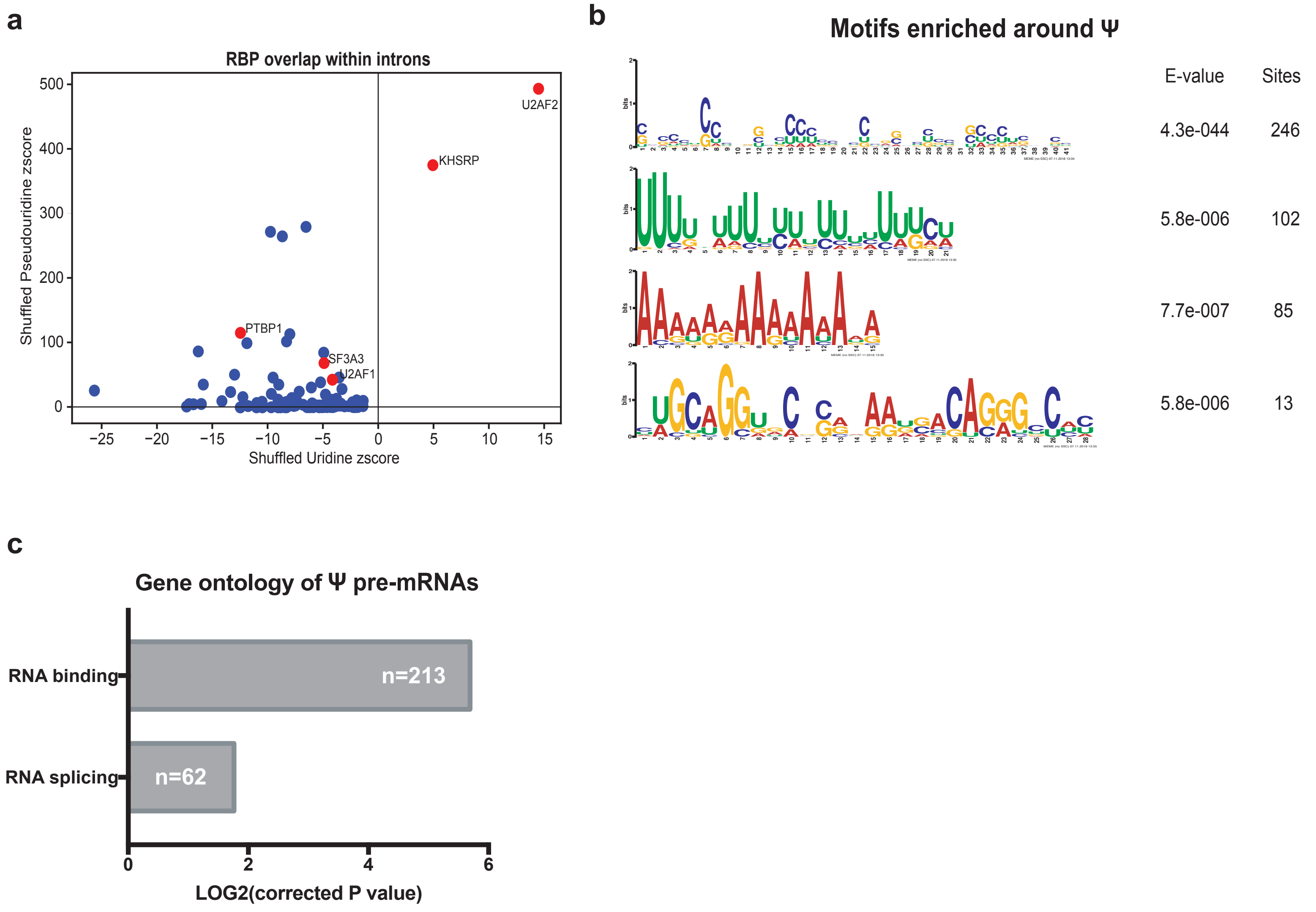

### Extended Data Figure 4

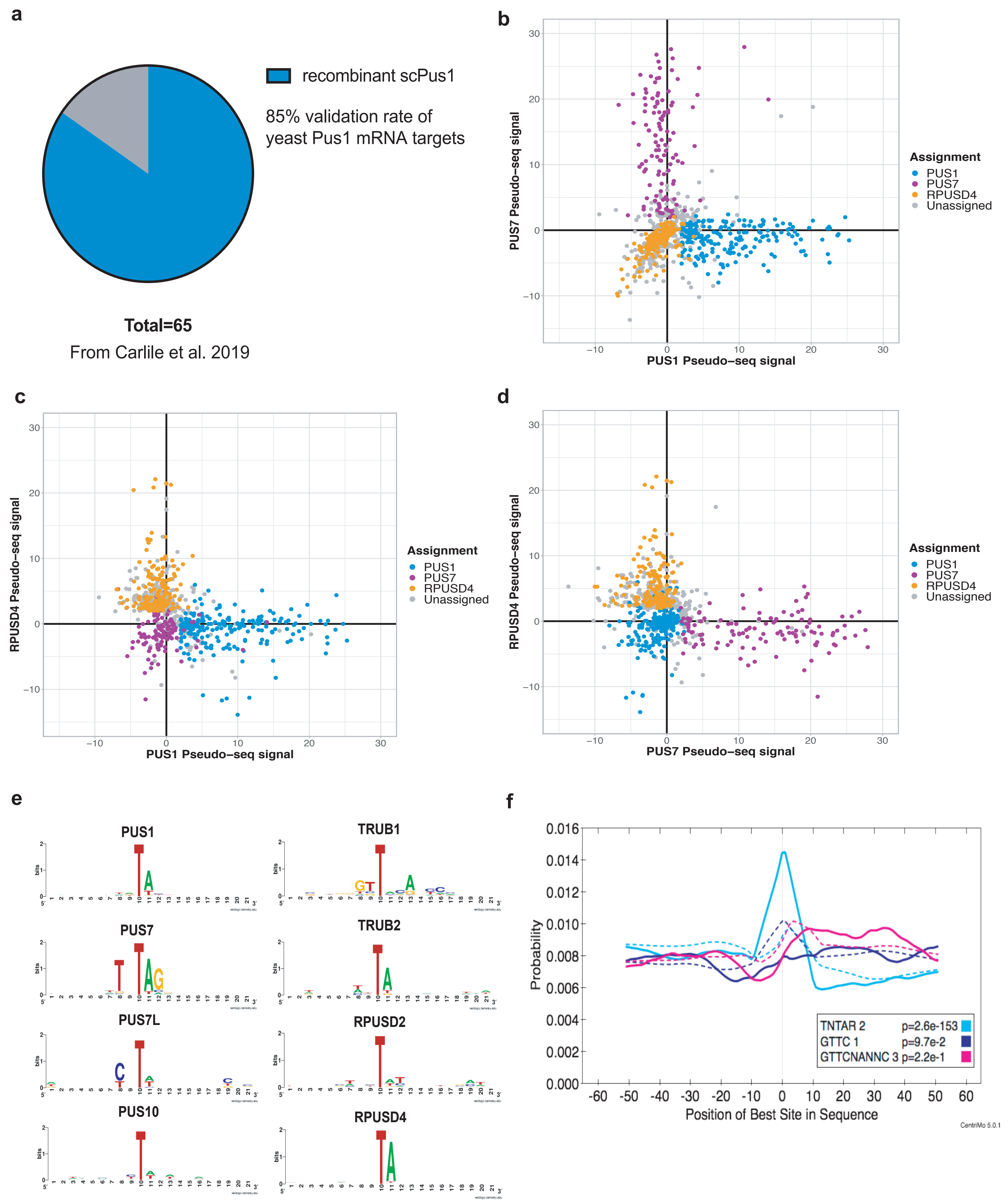

### Extended Data Figure 5

a

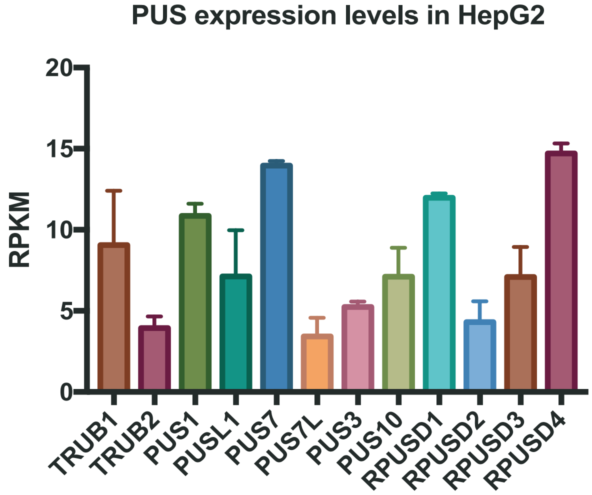


b

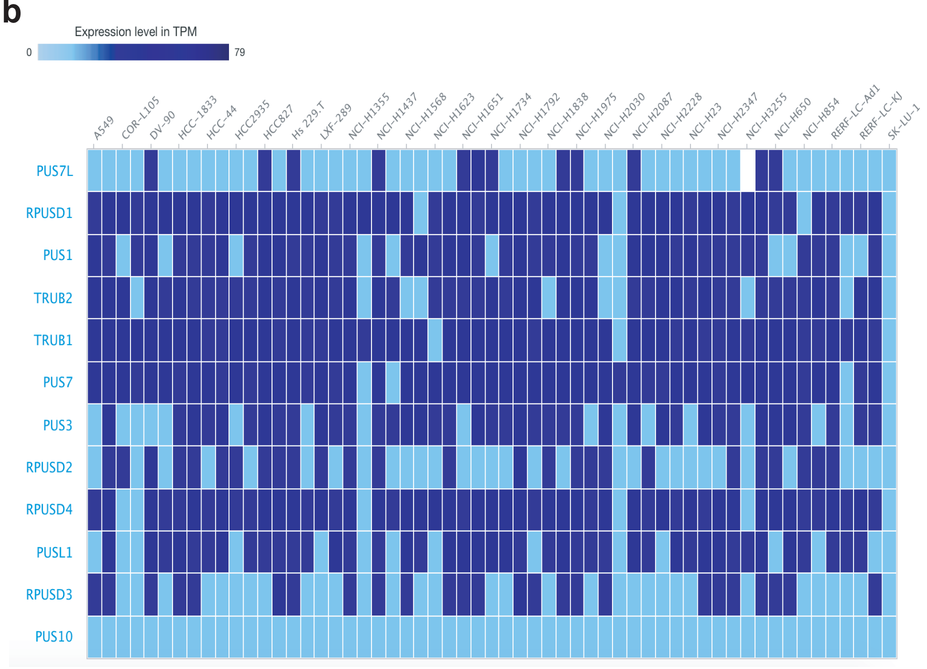


c

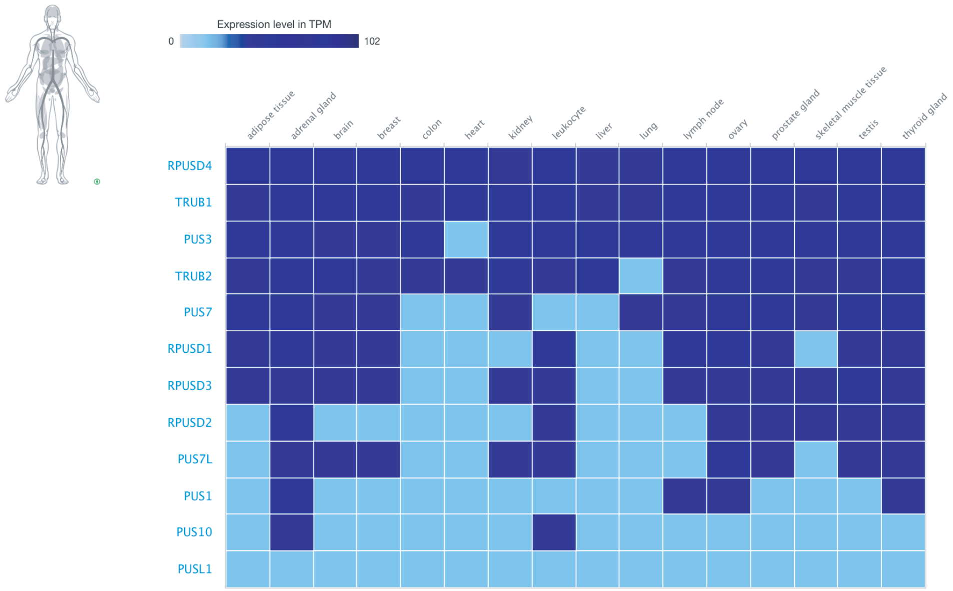

### Extended Data Figure 6

**a**

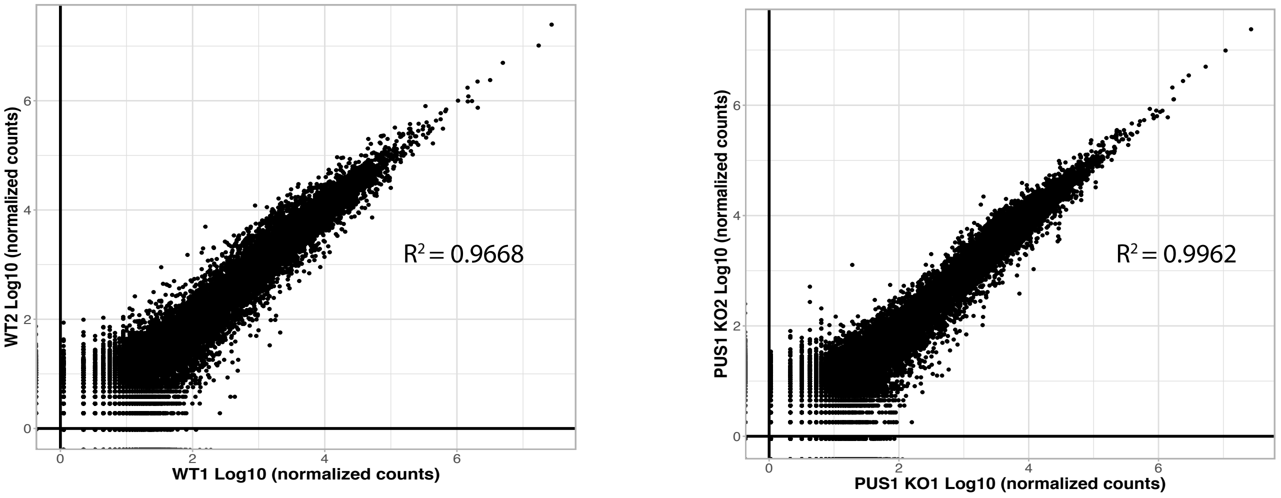


**b**

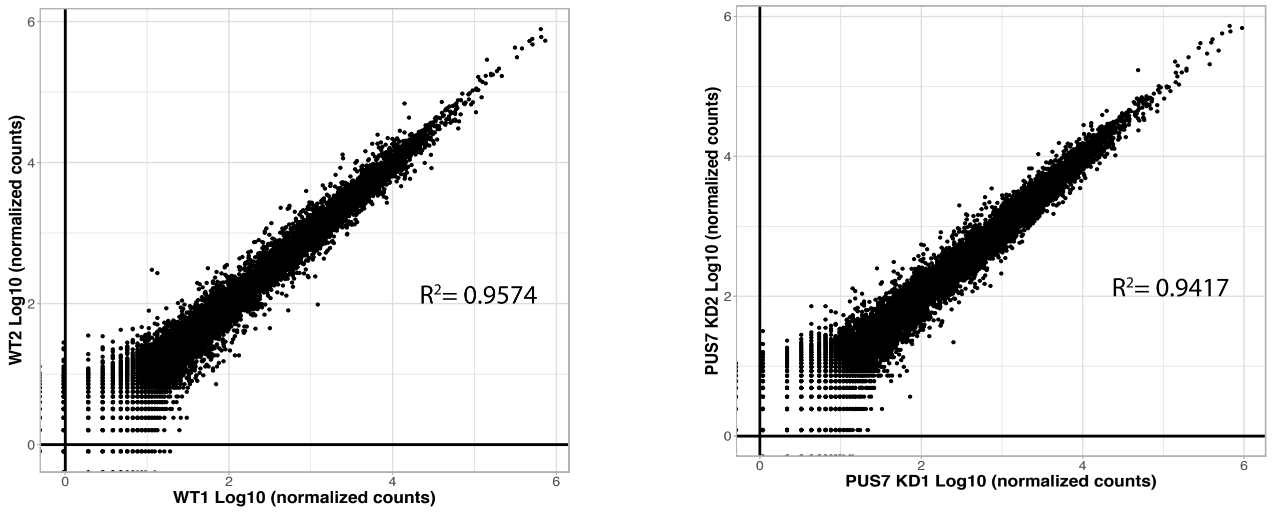


**c**

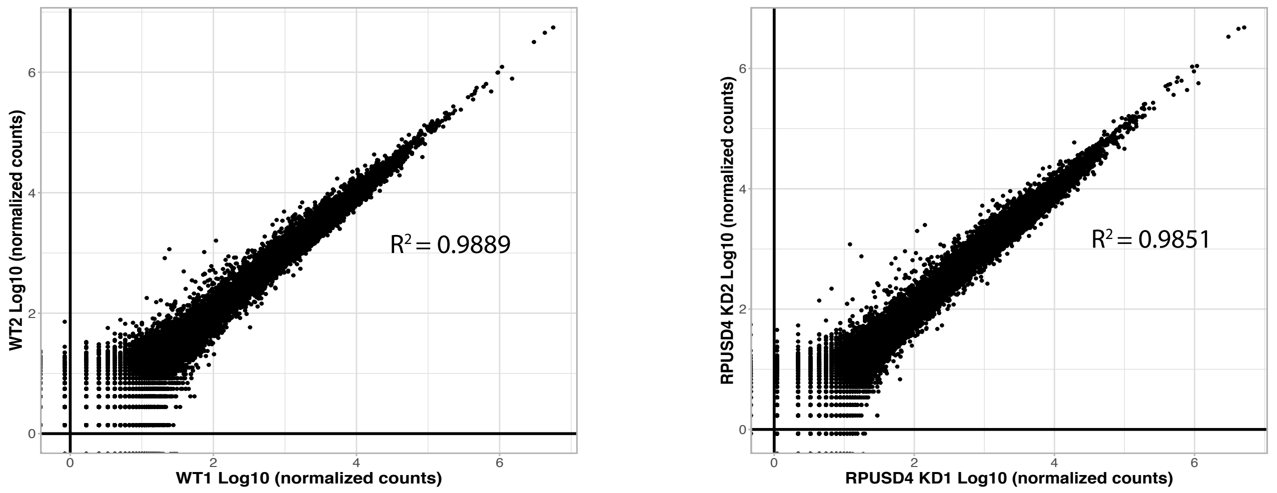

### Extended Data Figure 7

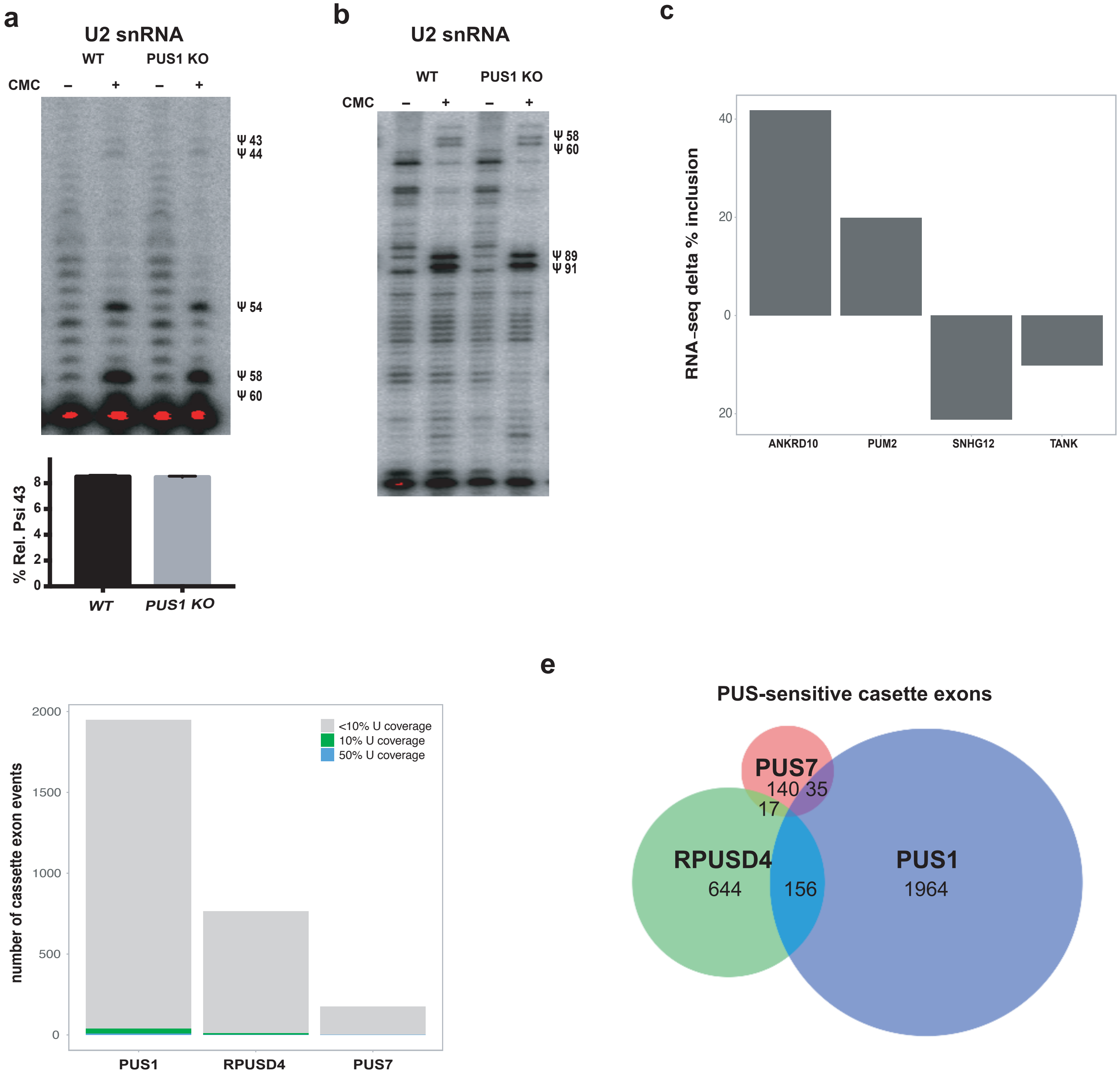

#### Extended Data Figure 8

a

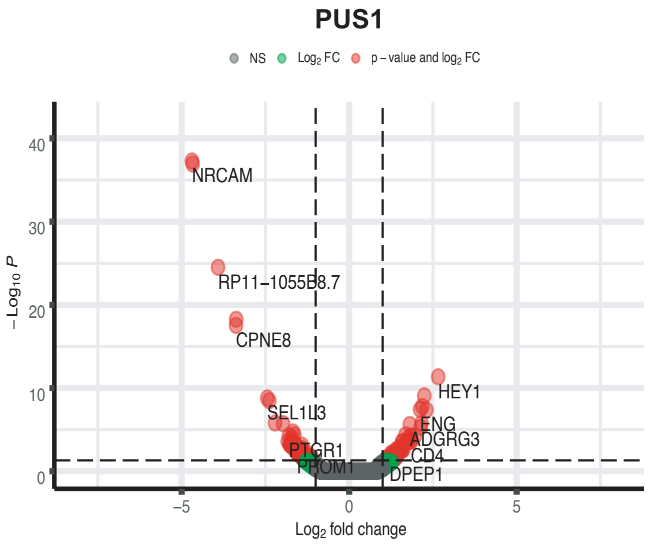


b

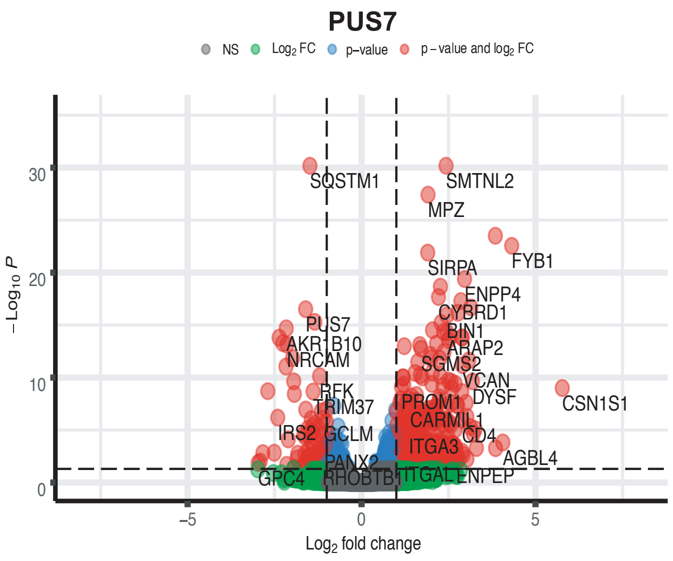


c

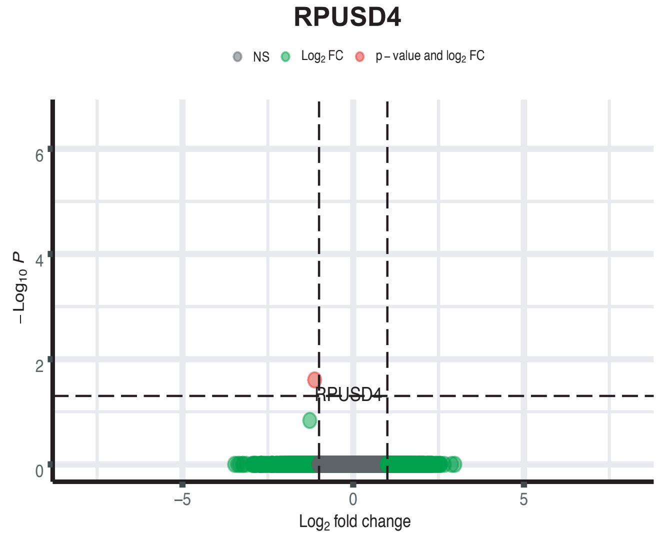

#### Extended Data Figure 9

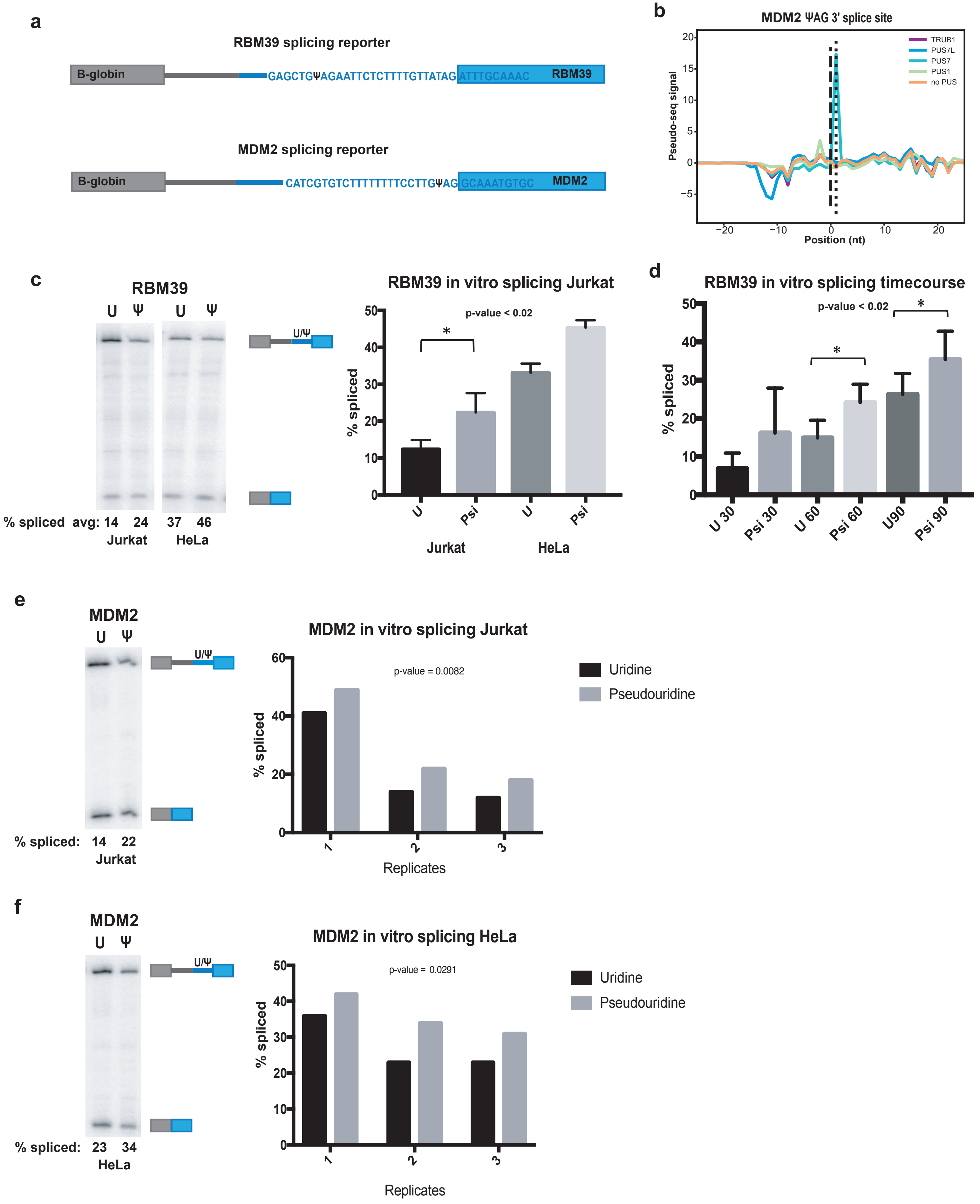
